# Cognition does not automatically influence perception: Evidence from neural encoding of colours belonging to different categories

**DOI:** 10.64898/2025.12.31.697186

**Authors:** Jasna Martinovic, Alexey A. Delov, Jana Tomastikova, Joel Martin, Galina V. Paramei, Yulia A. Griber

**Affiliations:** School of Philosophy, Psychology and Language Sciences, University of Edinburgh, EH8 9JZ, UK; School of Psychology, Liverpool Hope University, L16 9JD, UK; Department of Sociology and Philosophy, Smolensk State University, 214000, Russia

**Author notes:** Corresponding author: Prof Jasna Martinovic, 7 George Square (rest of address as above).

**Keywords:** colour, cognition, categorical perception, electrophysiology, Whorfian hypothesis

## Abstract

The firmest evidence in favour of models that posit early high-level influences of cognition on perception comes from electroencephalography (EEG). Enhanced early, pre-attentive processing of light and dark blue feature changes compared to light and dark green changes was reported in Greek speakers, who have two basic terms for ‘blue’ (*ghalazio/ble*). In the present three-experiment study, we systematically re-evaluate this evidence and test an alternative model that the early difference waves in the human EEG instead mainly reflect contrast adaptation phenomena. We use the same classical oddball paradigm presenting alternating standards and deviants that systematically differ in colour and/or luminance and chromatic contrast. We then calculate the visual mismatch negativity (vMMN), a putative index of pre-attentive feature change processing and predictive coding derived from EEG data, by subtracting the activity elicited by standards from that elicited by task-irrelevant deviants. Our experiments demonstrate the following: 1) vMMN is driven by contrast adaptation, being observable only in the presence of contrast differences between the stimuli and not reliably observed for categorically different hues equated in contrast; 2) there is no reliable difference between green- and blue-related difference waves in speakers (Russian) with two basic blue colour categories, the difference waves, again, being driven by contrast rather than their categorical content. Our findings are highly significant for the debate concerning the interface between perception and cognition, as the absence of early categorical effects speaks against models that predict Whorfian-type pre-attentive cognitive influences on perception.

**Significance statement:** The Whorfian hypothesis posits that basic language categories alter one’s perception of the world in a fundamental manner. Some of the most compelling evidence in favour of this hypothesis came from electrophysiological responses that indicated early differences between speakers of languages with different number of basic colour categories. The brain response taken as an indicator of these differences was considered to be a robust marker of early, pre-attentive processing and predictive coding. In the current multi-experiment study, we present evidence that early electrophysiological differences to colour reflect signatures of hue, saturation and luminance contrast and adaptation to this contrast, rather than of linguistic categories. This means that evidence in favour of early, pre-attentive categorical colour perception has been significantly eroded.

## Introduction

A key question in cognitive science is whether linguistic categories and concepts can directly shape perceptual processes, as proposed by the Whorfian hypothesis. The cognitive domain of colour categories presents a particularly salient natural experiment for evaluating putative Whorfian effects, since some languages differ in how they categorically partition the colour space(1–3). Furthermore, with its millisecond temporal resolution, electroencephalography (EEG) is a particularly informative tool for testing claims about early perceptual and cognitive processing. Thus, colour category research – both behavioural(3, 4) and electrophysiological(5, 6) – has played an important role in the ongoing debate on the extent of Whorfian effects on colour perception and the penetrability of perception by cognition more generally.

As discussed in the keynote dialogue of Ahissar and Scholl on the topic of cognitive penetrability of perception at the European Conference of Visual Perception in 2017, the most robust and direct evidence in favour of penetrability remains the event-related potential (ERP) study by Thierry and colleagues on Greek ‘blues’(5). ERPs represent EEG responses time-locked to an event of interest, averaged across multiple trials to remove non-systematic noise. In cognitive electrophysiology, different ERP components are taken as indices of different cognitive processes(7). The visual mismatch negativity (vMMN) is one such component – a negative deflection in the ERP difference wave peaking at about 150-240 ms when elicited by different colours(8). It is obtained by subtracting the event-related neural response driven by the repeated, standard stimuli from the event-related response to rare, deviant stimuli. The vMMN is assumed to index a mismatch between the built-up sensory representation of the standard stimulus and a rare deviant stimulus(9, 10). Thierry et al.(5) used a classical vMMN oddball paradigm and found greater vMMN resulting from light and dark blue series of standards and deviants compared to light and dark green series in Greek speakers, who have two basic terms for ‘blue’ (*ghalazio/ble*).

Thierry et al.’s study is considered to provide strong evidence in favour of models that posit high-level, top-down categorical effects on early, pre-attentive processing of colour(11) and goes against the more recent categorical facilitation model, which argues that effects of colour categories on perceptual processing are late and mediated by attention(12, 13). Thierry et al.’s findings on automatic categorical effects on early brain activity in the absence of a task that specifically recruits language also go against models that propose linguistic categories do not come into play in non-linguistic versions of very similar tasks(14). However, a recent critical review of EEG studies on colour categorisation between speakers of different languages identifies some major problems with this body of work and asks for a methodologically sound re-evaluation of the evidence base(15). These concerns are further compounded by specific issues with validity of early electrophysiological markers of visual feature-change detection and their robustness to contrast-adaptation effects(16).

Since classical oddball tasks involve streams of repeated stimuli (standards) interspersed by rare stimuli (deviants), with each standard stimulus repetition, there is a build-up of neural adaptation. In ERPs, adaptation manifests as repetition suppression(16), whose effects on the early, sensory parts of the EEG waveform are separable from expectation effects(17). Stimulus repetition led to a more negative waveform in the period of early visual evoked activity – reducing the amplitude of the P1 component and increasing the amplitude of the subsequent N1 component (17). Since vMMN overlaps in time with the P1 and N1, it is particularly vulnerable to repetition suppression. In the auditory domain, rapid and stimulus-specific adaptation has already been proposed as the underlying mechanism for a similar, standard-stimulus driven build-up in early positivity in evoked activity, related to echoic memory traces(18) and considered as the foundation of predictive coding(9).

After experimentally controlling for the effects of adaptation, the vMMN could not be reliably observed in response to orientation, luminance contrast, phase, and spatial frequency features of Gabor patches(16). Instead, an early deviant-related positivity (∼100 ms) was observed,which led to the proposal that, rather than the vMMN, this positive deflection might in fact be the neural marker of an early predictive processing of elementary visual feature values. The N1 component, which temporally coincides with the vMMN, exhibits both adaptation and response selectivity for colour (19). If we identify expectation violations, rather than simple repetition suppression effects with the underlying categorical perception processes that colour oddball tasks were intended to tap into, this poses a very concrete challenge to earlier EEG studies reliant on negative deflections such as the vMMN as an indicator of automatic, pre-attentive feature-change detection.

Considering the fundamental challenge to the validity of the vMMN as the index of early pre-attentive feature-change processing and the ongoing theoretical debates on the automaticity of categorical effects, it is important to re-evaluate Thierry et al.’s findings and provide more robust evidence for theories of colour cognition, and, more generally, for models of the perception/cognition interface. Here, we do that in three experiments, whose key aspects are depicted in Figure 1:

(1) A close replication of Thierry et al.’s(5) paradigm conducted, in place of Greek speakers, with Russian speakers, who, too, have two basic terms for ‘blue’ (*goluboj* ‘light blue’, *sinij* ‘dark blue’). The stimulus coordinates, oddball structure, and analysis approach closely followed the original study, while the language group differed.
(2) A conceptual extension with English speakers, with standards and deviants in the ‘cool’ and ‘warm’ regions of colour space, respectively. For English speakers, the ‘cool’ region includes two basic colour terms (BCTs) – *blue* and *green*, whereas the ‘warm’ region is lexically represented by four BCTs – *red, pink, yellow* and *brown*. Pairs of light and dark blue and light and dark green belong to the *blue* and *green* BCTs, respectively; in comparison, light and dark shades in the red region should be labelled by distinct BCTs *pink* and *red*, and light and dark shades in the yellow region should be labelled by distinct BCTs *yellow* and *brown*.
(3) A final experiment in which we test whether the vMMN can be reliably observed for feature changes along three perceptual dimensions of colour: hue, lightness, and saturation (colourfulness). If the vMMN is driven by a categorical distinction, it should be observed in all instances, as each block contains categorically distinct variations of ‘red’ and ‘green’. Alternatively, if the vMMN depends on a co-occurring difference in colour or luminance contrast, we should observe it only when contrast differences between ‘red’ and ‘green’ are introduced.

**Figure 1.**
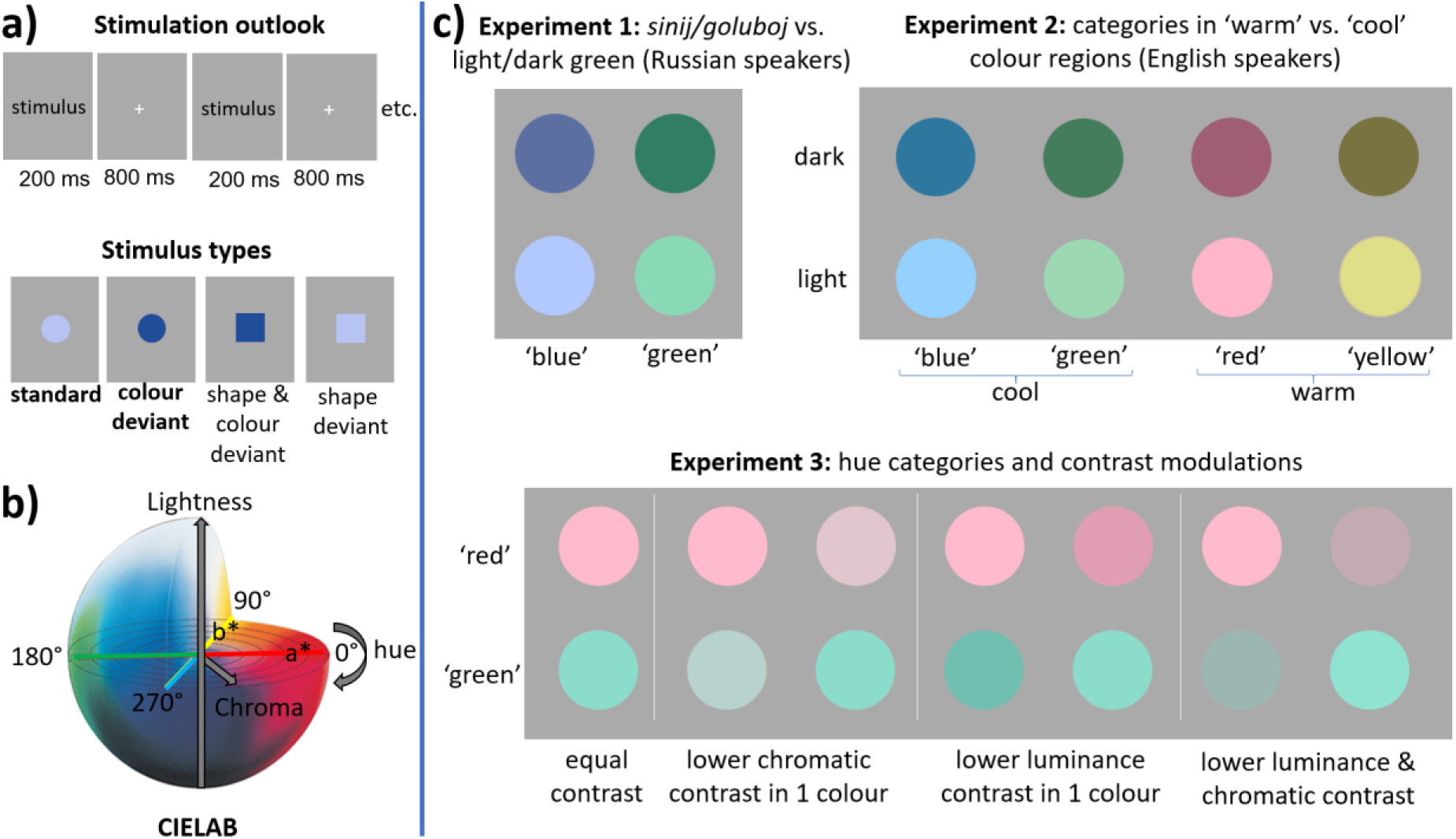
Experimental stimuli and the oddball paradigm. A) Stimuli are presented in pseudorandom sequences of 3-5 standard stimuli of circular shape, followed by an oddball that is either a deviant only in colour, and thus task-irrelevant, or is also a target (i.e., square) shape. The example stimulus types are taken from a block in which light blue is the standard colour and dark blue the deviant colour. B) CIELAB colour space, with lightness (L; range 0-100) as its vertical dimension, chroma (C; zero being achromatic) as the distance from the central lightness axis which defines the colourfulness of the stimulus, and hue as the angle of rotation on any of the horizontal planes within the space. Red and green are on the opposite ends of the 0°-180° axis, labelled as a*, while blue and yellow lie along the 90°-270° axis, labelled as b*. Stimulus colours from all experiments are plotted in CIELAB chromatic plane in Figure S1 in Supplementary Materials. C) Examples of stimulus appearance in each experiment. Appearance of the stimuli is approximated rather than exactly reproduced by converting all colours to sRGB whilst ensuring that RGB values are scaled proportionately to fit within the standard screen gamut. Note increased chromaticness and a slight shift towards purple in the blue sample of Experiment 1, which replicates the samples used by Thierry et al. (2009). Note also that in Experiment 3, red and green colours consistently belong to two different basic colour categories, but otherwise are either equal in chromatic or luminance contrast, i.e. different along one of these dimensions only, or different along both dimensions.

Foreshadowing our findings, we observe that EEG difference waves in the early time-window corresponding to the vMMN are driven by contrast adaptation, rather than by between-category differences. We fail to replicate Thierry et al.’s(5) observation of enhanced vMMN in speakers with two BCTs in the blue area of colour space, their study’s outcome likely stemming from the fact that the original observation had an effect size whose 95% confidence interval was very broad, indicating a less than precise estimation (μ_p_^2^ = 0.112, with a 95% CI of 0.004–0.273, estimated using the non-central F distribution(20)). Instead, we observe a series of robust, well-powered findings that consistently reflect effects of adaptation to luminance contrast, chromatic contrast, or both.

## Results

Figure 2 depicts ERPs and difference waves from our Experiment 1 with Russian speakers, side by side with replotted waveforms from the Thierry et al.(5) study, digitised from their EEG figure (21). Our ERPs elicited by blue and green colours follow similar patterns to those for the English sample in Thierry et al. The difference waves also appear similar, with the vMMN occurring in roughly the same time-window, followed by a positive deflection which results from a higher P300 response for task-irrelevant colour deviants. The P300 component reflects automatic attentional orienting (22), so it is unsurprising to see that it is higher for less frequent, task-irrelevant colour oddballs. The major difference in the two studies’ findings is that we observe a vMMN that appears to be lower in amplitude, so that evidence for robust differences against zero remains limited (see Figure 2a, bottom panel). This comparatively smaller vMMN also fails to exhibit any reliable differences between *sinij/goluboj* and light/dark green (BF_10_ = 0.301).

**Figure 2.**
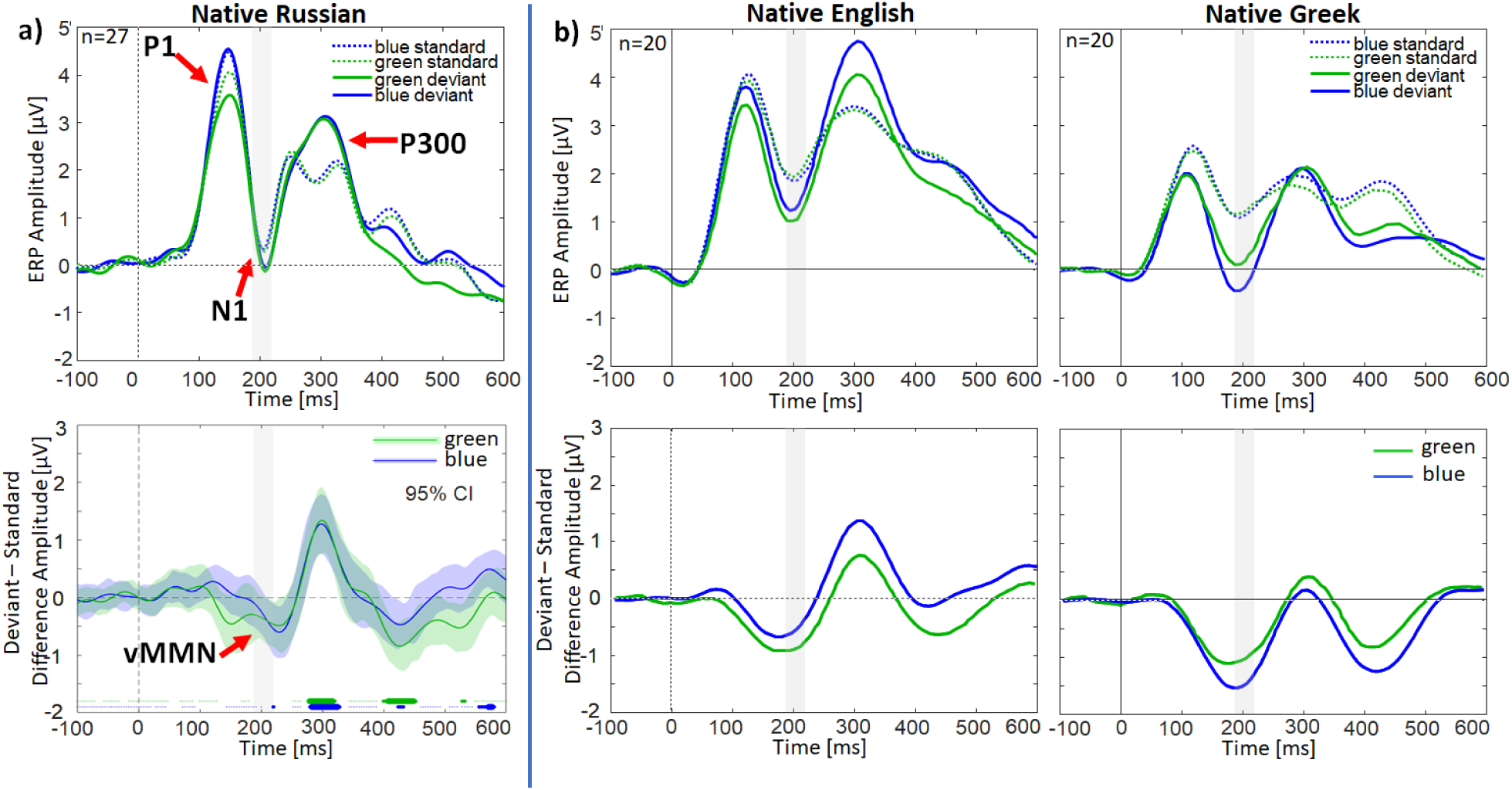
ERPs and difference waves from our sample of Russian speakers (panel a) and Thierry et al.’s samples of English and Greek speakers (panel b). Russian and Greek speakers are expected to exhibit a higher vMMN for blue due to two basic colour categories for dark and light blue, sinij/goluboj (Russian) or ble/ghalazio (Greek). However, the difference waves for ‘blue’ and ‘green’ in Russian speakers (panel a, bottom) are highly similar and small, with little evidence of a robust difference against zero. The top row depicts the grand-mean ERP waveforms elicited by green and blue standards and task-irrelevant deviants, which had identical colour coordinates across the two studies. EEG responses to task-relevant deviants (i.e., targets) are not shown, since they were explicitly attended and would, thus, not be useful in indexing automatic feature-change detection. The three most prominent ERP components in the stimulus-evoked response – P1, N1 and P300 – are marked by red arrows and labels in panel a. The bottom row depicts difference waves, created by subtraction of mean activity elicited by standards from that of identically coloured deviants. The vMMN is signposted by a red arrow and the label. In panel a’s depiction of the difference wave, Bayes factor (BF) values are indicated above the x -axis, with smallest dots indicating lack of significant differences from zero (BF < 0.33), medium-sized dots indicating moderate evidence for a significant difference (< 3 BF < 10), and large dots indicating strong evidence of a deflection in the difference wave (BF > 10). The time-window of the N1 peak in our study is highlighted by a light grey overlay across all panels, to facilitate comparisons.

In order to further understand the factors that drive EEG amplitudes, it is useful to depict ERPs and difference waves separately for dark and light blues and greens (Figure 3). This reveals that early evoked activity differs substantially between light and dark shades despite the fact that both colour sets had the same luminance contrast against the background (59%). In fact, lightness-driven differences are more substantial than between-hue differences: statistical analyses of ERP responses in the N1 time-window reveal a modulation only by lightness and deviance, which interact: amplitudes are more negative for light deviants (−0.96 µV, 95%CI [(−1.33), (−0.60)], t(182)=5.206, p<.001, 81% variance explained by the linear mixed effect model; for full model details, see Supplementary Materials 1). For lighter colours, we see moderate to extremely strong evidence of a robust negativity in the difference wave (light blue BF_10_ = 4.491, light green BF_10_ = 218.993), but there is anecdotal to strong evidence of absence of a vMMN-type negativity for darker colours (dark blue BF_10_ = 0.444; dark green BF_10_ = 0.250). Bayesian t-tests conducted across the time-course of the ERP (see bottom panel of Fig. 2a as well as Fig. 3b) corroborate these statistical analyses.

**Figure 3.**
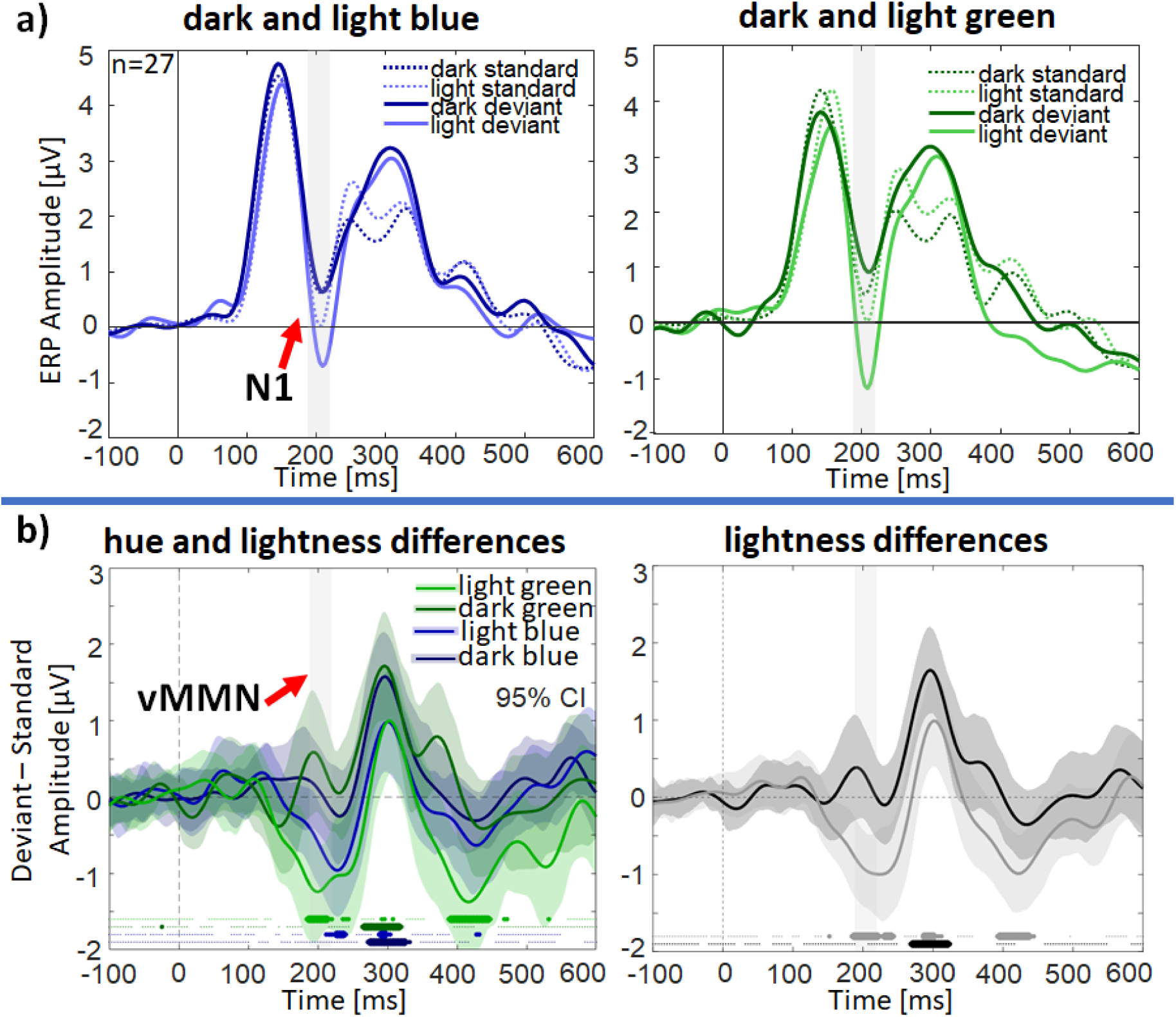
ERPs and difference waves from the experiment on Russian speakers, separated by lightness. (a) this panel depicts the grand-mean ERP waveforms elicited by light and dark standards and deviants for blue (left) and green (right panel). Note that during the windows characteristic of the N1 and vMMN components (shaded grey) it is the light standards and deviants that appear to elicit higher activity, and this is particularly the case with the light deviant, which would follow a set of repeated dark standards. (b) Difference waves, created by subtraction of light and dark standards from their respective deviants. The left panel depicts the difference waves separately for each hue/lightness combination. The right panel collapses across two hues to reveal differences by lightness alone. Bayes factor (BF) values are indicated above the x-axis, with smallest dots indicating lack of significant differences from zero (BF < 0.33), medium-sized dots indicating moderate evidence for a significant difference (3 < BF < 10) and large dots indicating strong evidence of a deflection in the difference wave (BF > 10). Note that averaging colours across lightness, which is depicted in Figure 2a, conceals that for dark greens and blues, there is no robust vMMN present, with the component only being notable for light colours.

In Experiment 2, conceptual replication with English speakers (Figure 4; for Bayesian stats across the difference wave timecourse, see Figure 4b), we fail to observe a reliable vMMN driven by ‘warm’ (BF_10_ = 0.134) or ‘cool’ colours (BF_10_ = 0.436), despite the fact that ‘warm’ colours are denoted by two BCTs across their light and dark variants (*pink/red* or *yellow/brown*), unlike ‘cool’ colours, light and dark green or blue, which are denoted by varied hyponyms or achromatic-modified terms (for names provided by our participants, see Supplementary Materials 3). In Experiment 2, we introduced a small difference (12%) in luminance contrast between dark (62%) and light (50%) shades of each colour. The luminance contrast difference is too small to alter categorical or relative (dark or light) belongingness, but is expected to elicit different degrees of adaptation of ON (light) and OFF (dark) luminance mechanisms(23). In line with this experimental manipulation, we observe difference wave modulations by luminance contrast, with a significant vMMN for lighter (BF_10_ = 2.340) but not darker colours (BF_10_ = 0.142) in the 170-220 ms time-window. Figure 4b reveals that this is due to a contrast-dependent shift in the latency of the vMMN component, with the vMMN appearing earlier for darker as opposed to lighter colours. For darker colours, the subsequent P300 deflection in the difference wave also appears earlier, resulting in the average amplitude being approximately zero in the N1/vMMN time-window. There is also an early positive component in the difference wave at ∼110 ms, coinciding with the P1 component of the ERP, which is robustly present for darker colours only (BF_10_=2.03; light colours BF_10_=0.21; see also Fig. 4b). The enhanced P1 for darker colours is in line with its higher contrast and with earlier and stronger luminance OFF responses(24), again, conforming with an explanation of early evoked differences as the signature of contrast and adaptation to it, rather than of categorical processing. This time, our statistical analyses of ERP amplitudes in the vMMN window do reveal a three-way interaction in the linear mixed effect model: there is a significant difference between standards and deviants only for light ‘cool’ colours (−0.762) µV, 95% CI [0.407,1.117]; t(519) = 4.210, p<.001; see Supplementary Materials 3 for more details). This replicates our observation of higher N1 amplitudes for light blue and light green in the 1^st^ experiment. Thus, instead of the predicted vMMN enhancement for ‘warm’ colours, we find the opposite pattern, although simulated power analysis(25) demonstrates we have only just enough power to observe this small effect (marginal R^2^ = 0.026, conditional R^2^ = 0.814; see Supplementary Materials 3 for further details).

**Figure 4.**
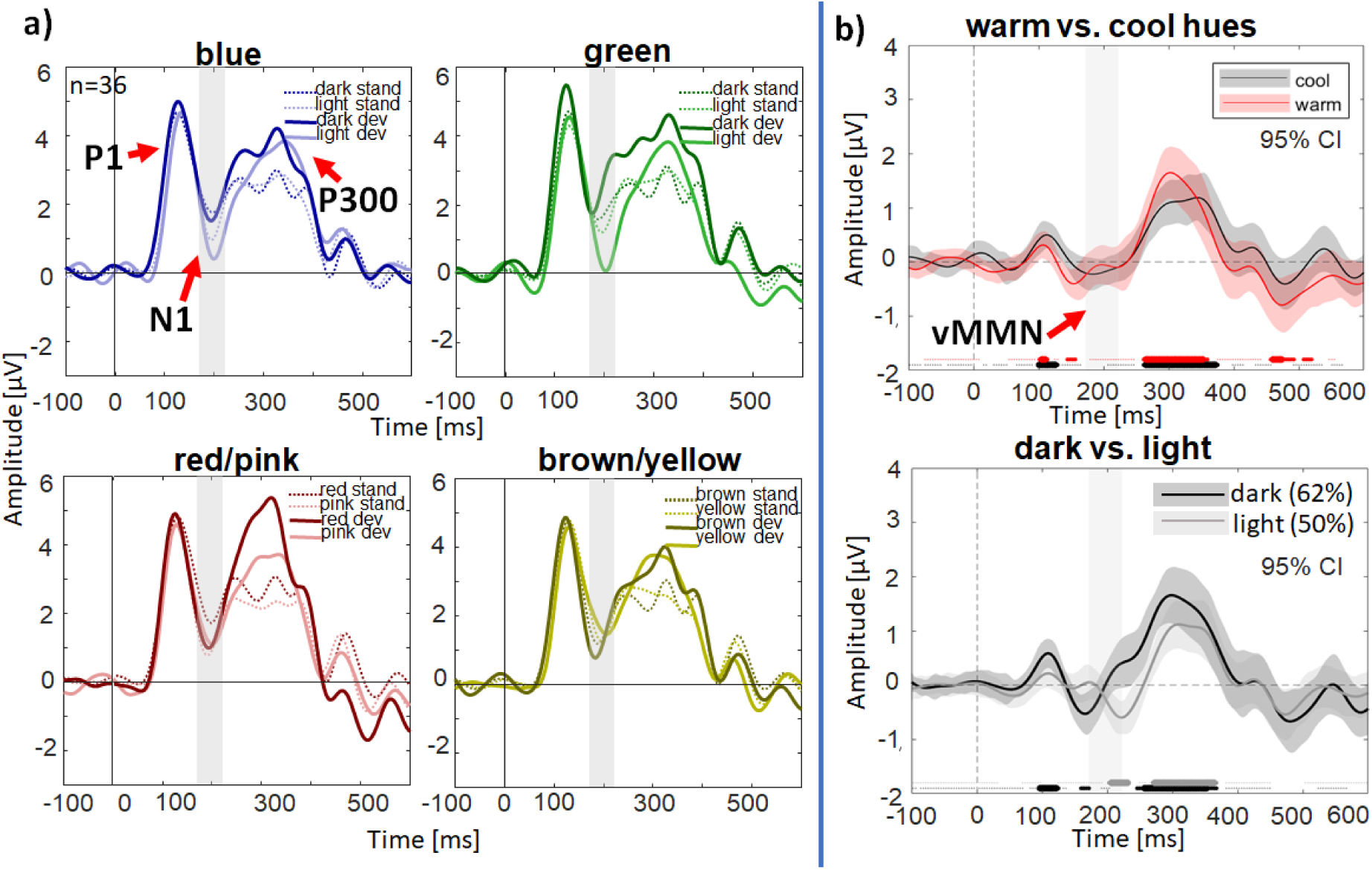
ERPs and difference waves from Experiment 2 with English speakers, who are expected to exhibit a higher vMMN for ‘warm’ hues due to multiple basic colour categories denoted in that region of colour space (i.e., red/pink, or yellow/brown), as opposed to less categorically and lexically distinguishable ‘cool’ region (green and blue). (a) This panel depicts the grand-mean waveforms elicited by light and dark standards and deviants for light/dark blue and light/dark green (top panels), and red/pink and brown/yellow (bottom panel). The overall waveforms are similar to those in Figures 2 and 3, with P1, N1 and P300 components being most prominent. Note, however, the apparent differences in the latency of the N1 response in the grey-shaded window between dark and light versions of a colour (most pronounced for green and yellow), with the N1 peak appearing earlier for darker shades. (b) Difference waves, created by subtraction of warm and cool (top) or light and dark (bottom panel) standards from their respective deviants. Bayes factor (BF) values are indicated above the x-axis, with smallest dots indicating lack of significant differences from zero (BF < 0.33), medium-sized dots indicating moderate evidence for a significant difference (3 < BF < 10) and large dots indicating strong evidence of a deflection in the difference wave (BF > 10). Two points are of note: (1) The difference wave for warm and cool hues does not exhibit a significant deflection in the N1 window, but there appears to be an earlier positive deflection at ∼110 ms, coinciding with the P1 window of the ERP; (2) As in Figure 2a, averaging hues across lightness levels conceals differences between dark and light colours. We see evidence for a positive deflection at ∼110 ms and an earlier subsequent vMMN for dark colours, which also happen to be higher in luminance contrast against the background. The effects of contrast on the vMMN for dark vs. light colours with an absence of a reliable effect of the category for warm vs. cool hues is, again, inconsistent with an interpretation of early evoked differences in the EEG as a signature of categorical processing. The time-window of the N1 and vMMN components is highlighted with a grey overlay.

Finally, to verify the observed effects of contrast on the vMMN, we conducted Experiment 3 in which we modulated hue, chromatic and luminance contrast systematically. Consistent with the findings from our first two experiments, vMMN responses were primarily driven by contrast-based properties (see Figure 5). Bayesian hypothesis testing revealed no reliable effect of hue on the vMMN (BF_10_ = 0.197). Changes in saturation (chromatic contrast) elicited asymmetrical responses, with a more negative deflection observed only for saturated deviants relative to standards (see Figure 5b), while desaturated colours showed no such effect, yielding a non-significant overall vMMN for saturation (BF_10_ = 0.238). In comparison, for luminance contrast (darker colours had 30% contrast, as opposed to 84% contrast for lighter colours), we found strong evidence supporting the presence of the vMMN (BF(d<0) = 10.059). This, again, suggests that luminance contrast modulates vMMN amplitude, consistent with its dependence on contrast-adaptation mechanisms rather than on categorical colour processing.

**Figure 5.**
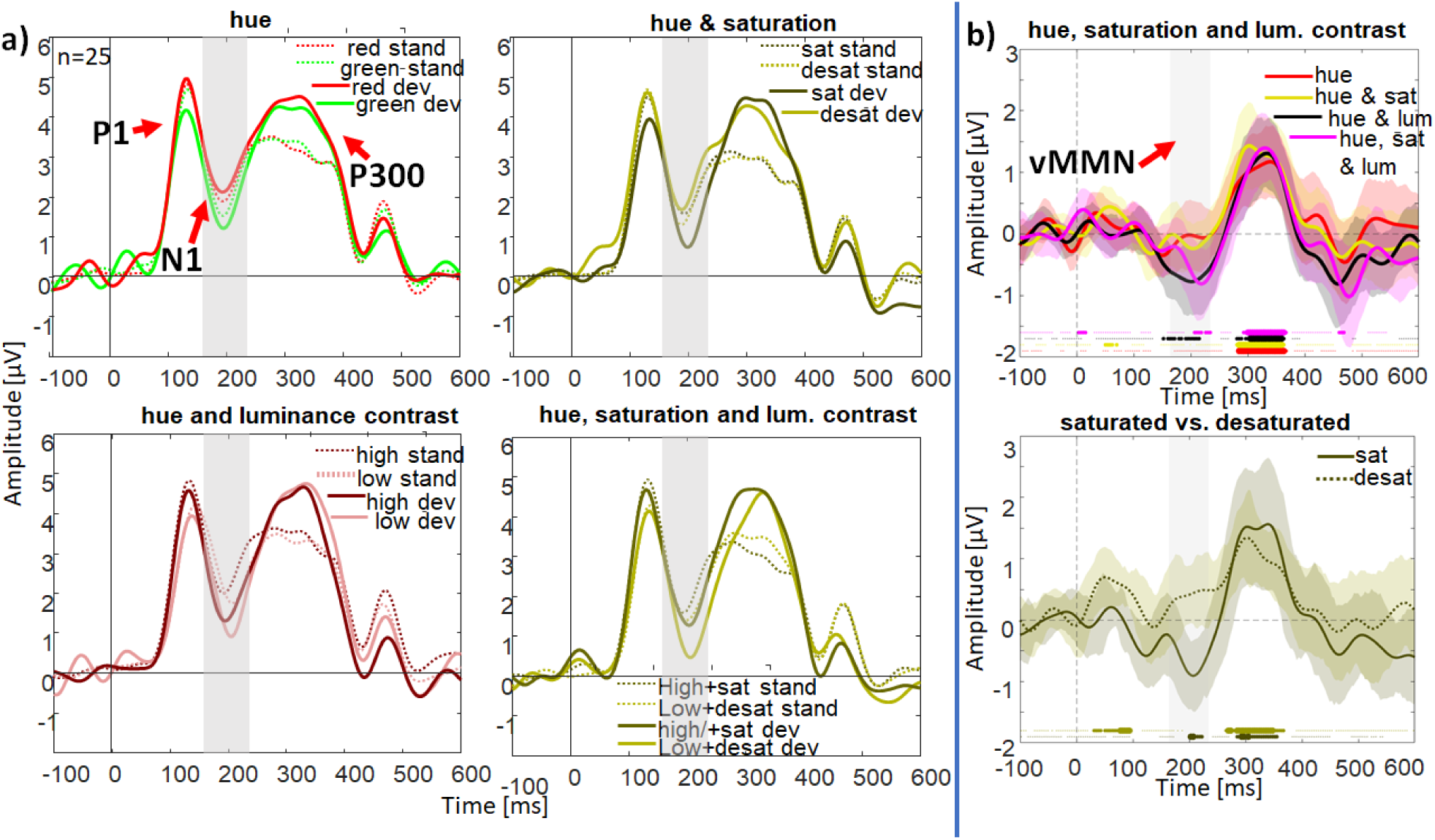
ERPs and difference waves from Experiment 3 examining the effects of hue, saturation and lightness. As stimuli are defined in CIE LCh space, L (lightness) is equivalent to luminance contrast and C (chroma, or colourfulness) corresponds to saturation, with higher chromatic contrast for more saturated stimuli. See also Figure S1 in Supplementary Materials for stimulus locations in both CIELAB and MacLeod-Boynton chromaticity diagram. (a) This panel depicts the grand-mean waveforms elicited by standards and deviants differing in only hue (red and green; top left), hue and saturation (top right), hue and lightness (bottom left) or all three dimensions (i.e., hue, saturation and lightness; bottom right). The overall waveforms are similar to those in Figures 2-4, with P1, N1 and P300 components. Note, however, the differences in P1 and N1 amplitudes and latencies, with P1 amplitude appearing to be higher and latency shorter for higher luminance contrasts. As expected, red also appears to elicit a higher P1 then green(26). (b) Difference waves, created by subtraction of deviants and standards in the four panels of Figure 5A (reflecting differences in hue alone, or hue together with saturation and/or lightness) or separated between saturated and desaturated hues (bottom panel). Bayes factor (BF) values are indicated above the x-axis, with smallest dots indicating lack of significant differences from zero (BF < 0.33), medium-sized dots indicating moderate evidence for a significant difference (3 < BF <10) and large dots indicating strong evidence of a deflection in the difference wave (BF > 10). Two points/aspects are of note: (1) The difference waves only exhibit a significant deflection in the N1 window when differences in luminance contrast are present. These differences are less robust across time for a combination of saturation and luminance contrast changes. (2) Averaging hues across saturation levels may conceal differences between saturated and desaturated colours. This could be the reason why combined effects of saturation and luminance contrast are less robust than for luminance contrast alone. The driver of these differences appears to be a more positive-going waveform for desaturated colours. In Supplementary Materials 4, we show a summation analysis of chromatic and luminance contrast combinations. As can be expected, this summation is not linear(27). Note that the time-window of the N1 and vMMN components is highlighted with a grey overlay in all subplots.

## Discussion

Our findings consistently and robustly demonstrate that colour categories do not automatically influence early neural markers of colour feature processing. In Experiment 1 we performed a close replication of Thierry et al.’s(5) widely cited study but with Russian rather than Greek participants, who also have two basic terms for light and dark ‘blue’. Although the two shades fall into separate ‘blue’ categories for Russian participants(28), we fail to observe robust categorical modulations. In fact, through two further experiments that systematically examine the factors that modulate EEG activity in the time-window of the early, perception-related processing of colour, we observe a series of consistent modulations by adaptation to luminance and colour contrast, with no modulations by basic colour categories. Our findings do not only challenge psychological models that posit an early, pre-attentive automatic influence of cognition on perception(11), but also raise questions that concern models of predictive coding(9), which propose some of the same visual contrast-driven EEG components as neural correlates of prediction.

How can we re-interpret the original finding of Thierry et al.(5) in light of our failed replication? Their study did not fully consider that the vMMN could be driven by factors other than categorical stimulus properties, e.g., lightness of a colour. If lightness was included in their statistical models, the authors would have observed that it appears to be the critical factor in eliciting the vMMN. The effects of lightness are not consistent with categorical effects and imply a more basic explanation for drivers of EEG differences. Our Experiment 3, manipulating hue, chromatic or luminance contrast, demonstrates that the vMMN is not robustly present for categorically distinct colours in red and green areas of colour space, which represent a highly salient categorical difference, but, rather, depends on chromatic and luminance contrast. Notably, while stimuli differing in luminance contrast elicit a robust vMMN, irrespective of whether the deviant or the standard has higher luminance contrast, for chromatic contrast the vMMN is elicited only for the higher-contrast deviant among lower-contrast standards. We know from normative studies that chromatic contrast is associated with negative deflections in the early ERP and lacks an initial positive peak(29, 30). The asymmetric effect of refractory activity following adaptation that we report here, to our knowledge, has not been previously reported. Asymmetric interactions between chromatic and luminance contrast are well known(31–33) and their manifestations in early electrophysiological markers of adaptation warrant further investigation.

If colour categories are not activated automatically, then the key determinant for their activation is likely to be the task that the participant is performing. It has been argued that this should be a task that involves language-derived colour concepts(14). This does pose challenges when probing categorical colour representation in pre-verbal infants or non-human animals, with recent studies opting for tasks reliant on recognition memory in pre-verbal infants (34) or working memory in macaques (35), since categorical biases are amplified when maintaining information in visual working memory (36). Future electrophysiology studies of colour categories should follow the Hillyard principle(7): using the same stimulus, they should examine the difference between perceptual tasks and tasks that require comparisons of a presented colour either to a previously seen sample, or to an internal, language-derived standard.

High susceptibility of the vMMN to contrast-adaptation effects is also very relevant for models of predictive coding. These models take MMN to be the signature of prediction across all sensory modalities(9). Male et al.(16) argue that the positive difference at ∼100 ms may be the signature of prediction for basic featural changes, instead of the vMMN. However, we demonstrate that even this positive deflection is contrast-related, since it is observed reliably only when stimuli vary in contrast and luminance polarity, and that its modulations are not independent of the subsequent modulations in the N1 window (see Figure S4). In fact, one could even argue that the oddball task, as employed by Thierry and colleagues(5), cannot elicit a genuine vMMN since the stimuli are all in the focus of attention. This would be corroborated by large P300 deflections that appear enhanced for deviants as opposed to standards (see Figures 2-5). Future studies will need to perform systematic tests of contrast vs. feature responsiveness at different retinal locations to establish whether foveal vMMN exists independently of contrast(37, 38).

In conclusion, our results reliably, consistently and systematically confirm the crucial role of contrast and contrast adaptation, rather than categorical basicness, in driving early event-related activity in the EEG in the first 200 ms of visual processing. This refutes some of the strongest evidence in favour of the models that propose a high degree of cognitive penetrability of perception(5, 11). Indeed, further studies of cognitive penetrability of perception could benefit from testing their predictions in the colour vision domain, where much is already known about the low-level drivers of visual evoked potentials(39), hence, potential low-level confounds are easier to predict and account for. This also applies to the search for neural markers of predictive coding. In fact, it would be highly interesting if in the visual domain they are built upon the probabilistic read-out of other, more basic and general biological mechanisms such as contrast adaptation(40). Having contrast adaptation at the root of prediction is not inconsistent with computational models of adaptation that involve ‘unaware’ read-outs(41). Indeed, examining the contributions of sensory adaptation to prediction anomalies would be a highly fruitful avenue for future research, of particular relevance for clinical applications of the model, which presume aberrant predictive coding in psychosis and often rely on MMN measures(42, 43).

## Methods

### Participants

**In Experiment 1,** 27 native Russian participants recruited from a university student population (10 male; all right-handed; mean age 23, range 19-25) completed experiment 1, which was a close replication of Thierry et al.(5). Three further participants were removed from the sample: one did not complete the experiment, another had an excessive target miss rate (56%), and still other produced insufficient artifact-free trials. All participants had normal or corrected-to-normal vision and showed normal colour vision assessed by the Farnsworth-Munsell 100 Hue test. Before participating, they gave written informed consent. The experiment was approved by the Ethics Committee of Smolensk State University.

Experiments 2 and 3 were conducted at the University of Edinburgh, with study protocols approved by the Psychology Ethics Committee. In the conceptual replication with English speakers (Experiment 2), 36 participants (6 male, 1 non-binary; 4 left-handed; mean age 25, range 19-36) were kept in the final sample, with 5 participants removed due to technical issues during the recording. Our final, pre-registered follow-up experiment on colour feature determinants of the vMMN (Experiment 3; https://osf.io/ymk9u/) included 25 participants (9 male, 2 left-handed, mean age 26, range 18-59) in the final sample, with 1 further participant removed due to an incomplete EEG session. All participants were recruited through posters around the university campus and word-of-mouth. They reported normal or corrected-to-normal visual acuity and had adequate colour vision, as assessed with the diagnostic plates from the City University test(44). Participants gave written informed consent and were reimbursed for their participation.

All three studies were conducted in line with the Declaration of Helsinki (1964).

### Apparatus, stimuli and procedure

Experimental stimuli and procedures were aligned as much as possible between the two experimental sites (Smolensk and Edinburgh), despite the reliance on different types of apparatus and EEG recording systems.

Experiment 1 (Smolensk) was run on an X-Rite™ Pantone® certified factory calibrated AERO 15 OLED 15.6-inch monitor with a screen resolution of 3840 x 2160 pixels at a refresh rate of 144 Hz viewed from a distance of 70 cm in a dark room with no other light source. Accuracy of colour reproduction on the monitor during the experiment was controlled with the help of the The Calibrite ColorChecker Display Pro device and software. The experiment was run using Psychophysics Toolbox Version 3 (PTB-3) software that extends Matlab (Mathworks, USA) with functions for research-grade stimulus presentation and response collection(45, 46). A VOROTEX K03 Red Switch (Vorotex, China) response pad with a simplified 3-key layout was used to collect behavioural data.

Experiments 2 and 3 (Edinburgh) were conducted using a calibrated Display++ screen (Cambridge Research Systems [CRS], UK) controlled by a ViSaGe visual stimulus generator (CRS, UK). The experiments were run using the CRS toolbox and CRS colour toolbox(47) for Matlab (Mathworks, USA). Measurements of monitor spectra were obtained using a SpectroCAL (CRS, UK) spectroradiometer. Participants were seated at a 70-cm distance from the display, which was positioned in a dark room with no other light source. They provided responses using a Cedrus-RB530 button box (Cedrus, USA).

Stimulus colours are depicted in Figure 1. Stimuli subtended 2° visual angle enabling use of 2° colour matching functions for calculating colour coordinates via the Optprop toolbox for Matlab(48). As in Thierry et al.(5), a grey background was set to CIE 1931 coordinates 0.3128, 0.3290, 26.2 cd/m^2^.

In Experiment 1, colours were identical to those used by Thierry et al. (CIE 1931, dark blue: 0.234, 0.230, 10.7 cd/m^2^, light blue: 0.259, 0.264, 41.5 cd/m^2^, dark green: 0.259, 0.397, 10.7 cd/m^2^, light green: 0.279, 0.377, 41.7 cd/m^2^). Luminance contrast against the background amounted to 59%, with LCh transformation of CIELAB coordinates for lightness (L), chroma (C) and hue (h) equal to L=39 for dark colours, L=71 for light colours, and C=28, h=281° for dark blue, C=28, h=278° for light blue, C=29, h=166° for dark green, and C=31, h=164° for light green. Transformation into CIELUV-derived LCh reveal a higher discrepancy between chroma of dark and light blues (46 vs. 41) and greens (39 vs. 31). The differences in chroma and hue between Munsell colours of different lightness, when transformed into CIELAB and CIELUV, are outcomes of the underlying transformations. Amongst other things, these calculations expand colourfulness at higher luminance levels(49). While we did not ask the participants in the EEG experiment to free-name the colour samples at the end of the experiment, we recruited a separate sample of 76 Russian speakers for this task. The dark blue stimulus was named sinij or tëmno-sinij ‘dark sinij’ by 81% participants, while the light blue stimulus was labelled goluboj by 32% of observers, with another 42% using different compounds containing goluboj or hyponyms of the goluboj category (for full naming data, see Supplementary Materials 2). These colour naming patterns are in line with previous work with Russian speakers(50).

In Experiment 2, colour coordinates were defined in CIELAB with these two aims in mind: 1) to ensure approximately equal saturation across hues and equal hue in the pairs varying in saturation only, by setting all colours to have equal chroma (C = 27) and hue angles (green h=155°, blue h=254°, red h=0° and yellow h=101°); 2) to introduce a small difference in luminance contrast (with luminance of 10.08 cd/m^2^ for dark and 39.34 cd/m^2^ for light colours, equal to 62% and 50% contrast against the background, respectively): whilst insufficient to alter the relational (light or dark) or categorical status (yellow or brown), such luminance contrast should still be sufficient to affect the EEG response systematically due to varying response properties for luminance ON and luminance OFF neurons(24). CIELAB was chosen as it accounts for differences in saturation more effectively than some other colour spaces(51).

We verified that colours indeed belonged to the same or different categories by asking participants to free-name all colour samples at the end of the experiment. Colour circles were presented on the screen in a randomised order; we report frequency and percentage of basic colour terms assigned to them, as well as of hyponyms, modified, compounded, or ‘fancy’ terms that could be allocated to those same categories (e.g., periwinkle for light blue; see Supplementary Materials 3 for the full list of names). Light and dark blue stimuli were named blue by 21 (58%) and 29 (81%) participants, respectively, with the remaining participants generally using non-basic terms that would fall under light or dark blue. A similar picture emerged for light and dark green, which were named green by 16 (44%) and 28 (78%) participants, respectively. Pink was offered by 27 (75%) of observers, while in naming of the red sample both pink (10 participants, 28%) and red (8 participants, 22%) were prominent (for similar patterns of pink-naming, see Rosenthal et al. (52)). Yellow was elicited in 30 (79%) participants, while naming of the brown stimulus was split across two basic terms, with 11 participants (31%) opting for green and the same number opting for brown. An increase of greenness in yellow colours with reduction of luminance was to be expected(53).

In Experiment 3, we selected our colours so as to manipulate hue, saturation and luminance contrast selectively – either in isolation or in combination with each other – to test the effects observed in the first two experiments in a principled manner. This experiment was preregistered prior to data collection, meaning that the hypothesis, design and analysis plan were publicly specified on the Open Science Framework (OSF; https://osf.io/ymk9u/). Colours – red and green – were defined in CIELAB LCh coordinates, which allowed us to systematically vary perceptual properties across blocks. Red was always set to h=0° and green to h=180° in CIE LCh. The experiment consisted of 16 blocks, organised into 8 block pairs, with standard and deviant roles reversed within each pair between red and green. Blocks 1-2 and 3-4 had stimuli of fixed lightness and chroma (L=75, C=27 for both red and green). In blocks 5-6 and 7-8, both hue and chroma were manipulated: in blocks 5-6, green was desaturated (L=75, C=27 for red and L=75, C=10 for green), while in blocks 7-8 red was desaturated (L=75, C=10 for red and L=75, C=27 for green). Blocks 9–10 and 11–12 manipulated hue in combination with luminance contrast, lowering lightness while keeping chroma constant (L = 75, C = 27, h = 0° vs. L = 65, C = 27, h = 180°; and L = 65, C = 27, h = 0° vs. L = 75, C = 27, h = 180° respectively). The luminance contrast manipulation amounted to 30% contrast for the darker colours, as opposed to 84% contrast for the lighter colours. Finally, blocks 13–14 and 15–16 introduced combined changes in all three dimensions—hue, chroma, and lightness. In blocks 13–14, both chroma and lightness were reduced for one hue (L = 75, C = 27, h = 0° vs. L = 65, C = 10, h = 180°), and in blocks 15–16, the asymmetry was reversed (L = 65, C = 10, h = 0° vs. L = 75, C = 27, h = 180°). Each block comprised 225 trials, of which 45 were deviant (either by colour, or shape, or combination of both).

Our experimental procedure followed closely the template provided by Thierry et al.(5). Stimuli were presented for 200 ms with an interstimulus interval of 800 ms. Streams of 3, 4 or 5 standard stimuli were followed by a randomly selected oddball stimulus resulting in stimulus streams with 6.67% oddballs of each type (shape only oddball, luminance and shape oddball, luminance only oddball). Participants were instructed to respond to shape oddballs (i.e., squares) and ignore circles. The stream contained the following total numbers of stimuli: 2700 in four blocks of 675 trials, which included 540 standards and 135 deviants (45 of each type) in Experiment 1; 3600 in eight blocks in Experiment 2, with 450 standards and 90 deviants of each colour (30 of each type; thus 60 non-target deviants, when collapsed across ‘warm’ and ‘cool’ hues) in Experiment 2; and 3600 in Experiment 3, separated into 16 blocks. Block order was randomised across participants. One participant from Experiment 1 was excluded due to low hit rate (<75%). Participants remaining in the sample performed the task with a high degree of accuracy (Experiment 1: 86.0% correct rejections, 0.7% false alarms, 13.0% hits, 0.3% misses, reported as a percentage out of a total of 100%, with average reaction time (RT) and standard deviation 600 ± 115 ms; Experiment 2: 86.5% correct rejections, 0.2% false alarms, 12.9% hits, 0.4% misses, RT 423 ± 95 ms; Experiment 3: 86.49% correct rejections, 0.18% false alarms, 13.06% of hits and 0.27% misses, RT 444 ± 98 ms).

### EEG recording and pre-processing

In Smolensk (Experiment 1), electrophysiological data were recorded continuously using an eego mylab 64-channel amplifier and waveguard original EEG caps, with 64 Ag/AgCl electrodes placed according to the extended 10–20 system. Data were sampled at 250 Hz. Bipolar horizontal (HEOG) and vertical (VEOG) electrooculograms were recorded simultaneously to monitor eye movements. Impedance of all electrodes was kept below 10 kΩ. Stim Tracker (Cedrus, San Pedro, CA, USA) was used to synchronise stimuli with the EEG data, bypass delays associated with operating systems and accurately record true onset time of the stimuli. In Edinburgh (Experiments 2 and 3), continuous EEG was recorded using a 64-channel BioSemi Active-Two (Biosemi, Netherlands) amplifier system, digitising the signal at a 1024 Hz sampling rate. 64 Ag-AgCl electrodes were mounted in an elastic, tight-fitting cap. In addition, we also recorded HEOGs and VEOGs, and two mastoid references. Stimulus triggers were sent from the Display++ device, synchronised to stimulus onset using Visage Framestore cycling functions.

Data from both sites were analysed in Matlab (Mathworks, US) using the EEGLAB toolbox(54). FASTER(55) and ADJUST(56) toolboxes were used for artifact rejection and correction. Initially the data were average-referenced and filtered using a 40 Hz low-pass and 0.1 Hz high-pass Hamming windowed sinc Finite Impulse Response (FIR) filters, as implemented in EEGlab(54). Recordings were then separated into epochs representing individual trials, 1000 ms in duration. This consisted of a 200-ms period before the stimulus onset and 800 ms thereafter. We also applied a baseline correction between −100 ms and 0 ms. Contaminated trials were identified and rejected using the FASTER toolbox. This is done by characterising global channel properties using statistical parameters of the data and applying a metric of ±3 z-score as a marker for contaminated data. Independent component analysis (ICA) was then performed using the default runica algorithm. The ADJUST toolbox was used to identify specific components driven by blinks, eye movements and local discontinuities. These were removed from the data. Following artifact rejection, FASTER was used to identify any contaminated channels and interpolate them from neighbouring channels. A threshold of ±3 was applied to the F-score parameter to define these outliers, which were removed and interpolated. A manual check was then performed on the data to verify quality of the artifact rejection procedure. In Experiment 1, percentage of rejected trials ranged between 1% and 5%; in Experiment 2, this was between 1% and 7%, and in Experiment 3 between 1% and 5%. As in Thierry et al.(5), the vMMN was quantified over parieto-occipital electrodes: in Experiment 1, these were O1, O2, Oz, PO7 and PO8, while in Experiments 2 and 3, Iz was also available in the electrode montage and, thus, added to this electrode cluster. Time-windows for statistical analyses were selected to encompass the N1 peak from the grand-mean ERP plots: 188-220 ms in Experiment 1, 172-223 ms in Experiment 2, and 164-234 ms in Experiment 3. The slight differences in the time-windows stem from the need to encompass the N1 component of the ERP, which varied in latency along expected lines in those experiments that manipulated contrast(30, 57).

### Statistical Data analysis

We fitted linear mixed effects models (LMEMs) to our data, with random by-participant intercepts to model overall differences in amplitude between observers and fixed effects of deviancy, hue and lightness (in Experiment 3, there was an additional fixed effect of saturation) to model our experimental manipulations, with simple coding of effects. To evaluate whether the ERP difference wave manifested a reliable difference from zero, we used Bayesian t-tests(46). Other statistical analyses were implemented in R 4.3.1(58) using the following packages: lmerTest 3.1-3(59), emmeans 1.10(60), DHARMa 0.4.6(61), simr 1.0.7(62), and BayesFactor 0.9.12(63). For further details of statistical analysis, see Supplementary Materials 2-4, where we also report statistical sensitivity/power analyses and all the properties of best fitting models.

## Acknowledgements

We thank Corinna Haenschel for helpful comments. Tabitha James, Emma Murray, Ruicong Yuan and Jiaheng Ou collected the data for Experiment 2 (Edinburgh). We are grateful to Thierry et al. (2009) authors Benjamin Dering and Jan-Rouke Kuipers for communicating with us about the percentage of oddballs in their study (6.67% per oddball condition, rather than 10% as reported in the original paper). Last but not least, we acknowledge funding from the Royal Society International Exchange Scheme and the Experimental Psychology Society undergraduate bursary scheme.

## Supplementary Materials

### Supplementary Material 1 – Stimulus locations in the CIELAB space and Macleod-Boynton chromaticity diagram across all three experiments

**Figure S1.**
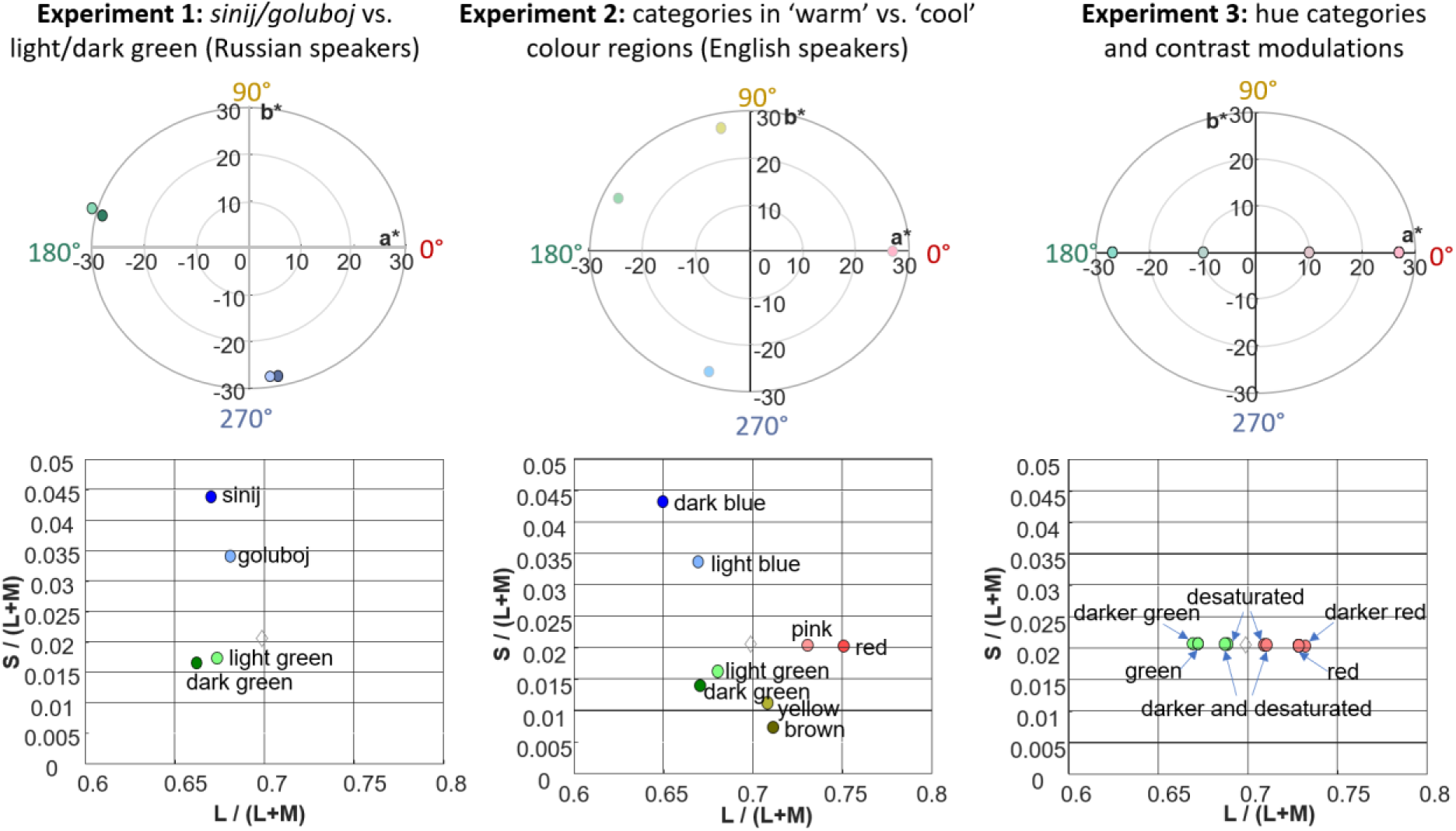
Top panel: Colour coordinates in the a*b* colour plane of the CIELAB space. For a full depiction of the CIELAB space, see Figure 1. a* captures the red (0°) – green (180°) dimension, while b* captures the blue (270°) - yellow (90°) dimension of hue. Distance from the centre of the space is equivalent to chroma. Bottom panel: Colour positions in the MacLeod-Boynton (MB) chromaticity diagram, whose axes depict L-cone (x) and S-cone (y) excitations, relative to the sum of the L and M-cone excitations. Note that green and red colours in Experiment 3 occupy approximately opposite locations relative to the background on the L axis. In CIELAB, our background’s chromaticity coincides with the centre of the space and in the MB diagram it is denoted with the grey diamond.

### Supplementary Material 2 – Effect of additional basic colour categories in the blue area of colour space (Experiment 1)

#### Best fitting linear mixed effect model

Fixed effects of deviancy (standard – reference level, or deviant) and lightness (dark – reference level, or light) and random by-participant intercepts only.

Model: amplitude ∼ deviancy*lightness + 1|participant

| Amplitude (μV) |  |  |  |
| --- | --- | --- | --- |
| Predictors | Estimates | 95% CI | P |
| (Intercept) | -0.12 | [(-0.86), 0.61] | 0.742 |
| Deviancy | -0.49 | [(-0.75),(-0.23)] | **<0.001** |
| Lightness | -0.81 | [(-1.07),(-0.56)] | **<0.001** |
| Deviancy × Lightness | -0.95 | [(-1.46),(-0.43)] | **0.001** |
| Random Effects |  |  |  |
| $\sigma^2$ | 0.87 | | |
| $\tau_{00}$ participant | 3.51 | | |
| ICC | 0.80 |  |  |
| $N_{\text{participant}}$ | 26 | | |
| Observations | 205 |  |  |
| Marginal $R^2$ / Conditional $R^2$ | 0.061 / 0.814 | | |

Prior to statistical analyses, we rejected any datapoints that were outliers: this resulted in rejection of 11 datapoints, which included all the datapoints from one participant (as can be seen from N and Observation rows in the table above).

Assumption checks were performed using the DHARMa R package and confirmed that the data were appropriate for the fitted model. We also verified that the predictors were not correlated by fitting an additive model and evaluating it with the *check_collinearity* function from the performance R package.

Initially, we fitted a model with the full set of fixed effects (deviancy*colour*lightness), but the step function from lmerTest R package indicated that colour (with two levels: blue and green) did not contribute significantly to the model, individually or in interaction with the other two factors:

Deviancy:Colour:Lightness, F(1, 171.98) = 3.4135, p = 0.066
Deviancy:Colour, F(1, 172.98) = 0.0339, p = 0.854
Colour:Lightness, F(1, 173.98) = 0.631, p = 0.428
Colour, F(1, 174.98) = 0.0436, p = 0.835

On the other hand, deviancy and lightness interacted significantly and therefore could not be removed from the model (F(1,175.98)=12.963, p<.001).

#### Power analysis

Sensitivity analyses were performed by resampling the number of participants (1000 simulations) and evaluating the observed power for obtaining the statistically significant fixed effects in the best fitting model, using the R package *simr*. Sufficient power is reached within 16 participants for the interaction of deviancy and lightness (see Figure S2, plotting simulated power on the y-axis against number of participants on the x-axis). We also evaluated the power to detect the observed effects if they were reduced by 15%, as advised by Kumle and colleagues(25)), and found that this was 91.70% (95% CI [89.81%,93.34%].

**Figure S2.**
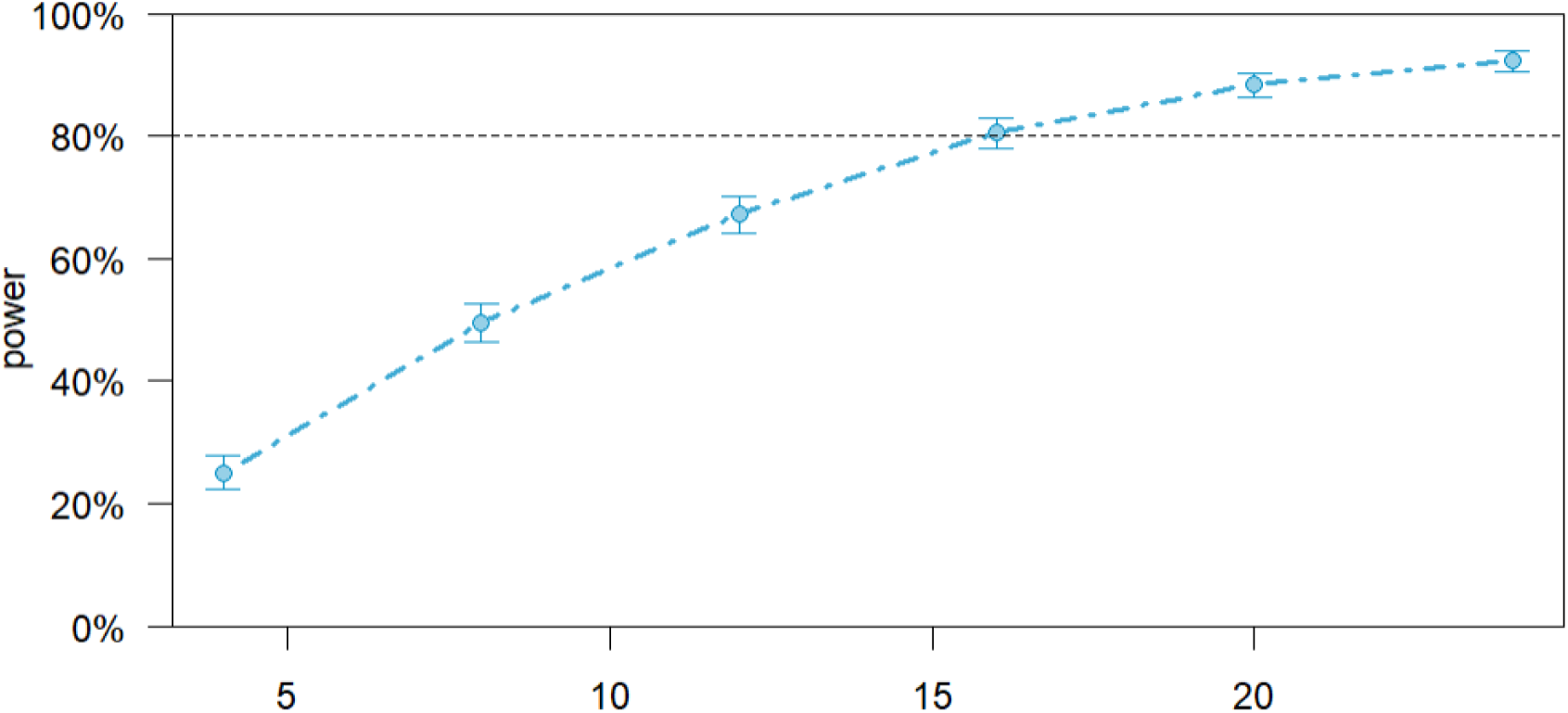
Power analysis for Experiment 1: deviancy by lightness interaction.

#### Free colour naming of colour circles by Russian participants

We verified that colours indeed belonged to the same or different ‘blue’ categories by asking 76 native Russian speakers (25 male; all right-handed, mean age 22, range 19-25) to free-name the four colour samples in a separate naming experiment, approved by the Ethics Committee of Smolensk State University. Colour circles were presented on the screen in a randomised order.

In Tables S1-4 show frequency (F) and relative frequency (%F) of colour terms assigned to each stimulus. Russian colour names were transliterated into Latin script using the free Online Transliterator (https://translit.cc). English glosses followed Frumkina and Mikhejev (1996).

We obtained the largest numbers of different colour names (basic colour terms, as well as hyponyms, modified, compounded, and ‘fancy’ terms) for the light blue (N=21) and light green (N=17) stimuli. The sets of colour names obtained for dark stimuli were more consistent and included 14 terms for dark blue and 13 for dark green.

Light green stimulus was named *svetlo-zelënyj* ‘light green’ or *zelënyj* ‘green’ by 61% participants, with the remaining participants generally using non-basic terms and compounds that would fall under light green (*mâtnyj* ‘mint’, *salatovyj* ‘lettuce-coloured’, *zelëno-birûzovyj* ‘green turquoise’, *bledno-birûzovyj* ‘pale turquoise’, *bledno-izumrudnyj* ‘pale emerald’, *bledno-zelënyj* ‘pale green’, *cvet morskoj volny* ‘colour of sea wave’, *izumrudnyj* ‘emerald’, *malaxitovyj* ‘malachite’, *svetlo-birûzovyj* ‘light turquoise’, *svetlo-izumrudnyj* ‘light emerald’, *svetlo-malaxitovyj* ‘light malachite’, *svetlo-salatovyj* ‘light-lettuce-coloured’, *tëmno-mâtnyj* ‘dark mint’, *xaki* ‘khaki’) (Table S1).

**Table S1.**
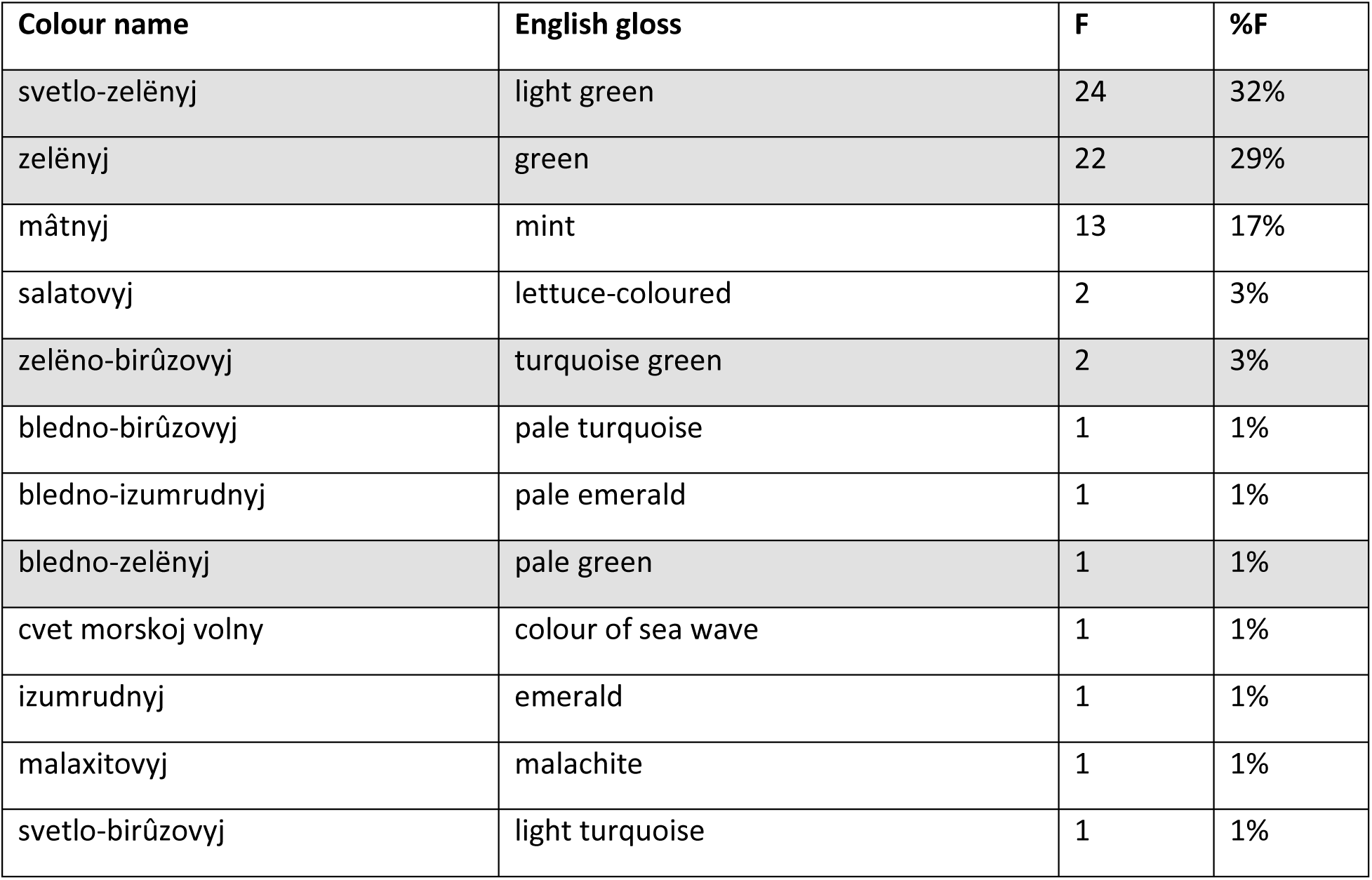
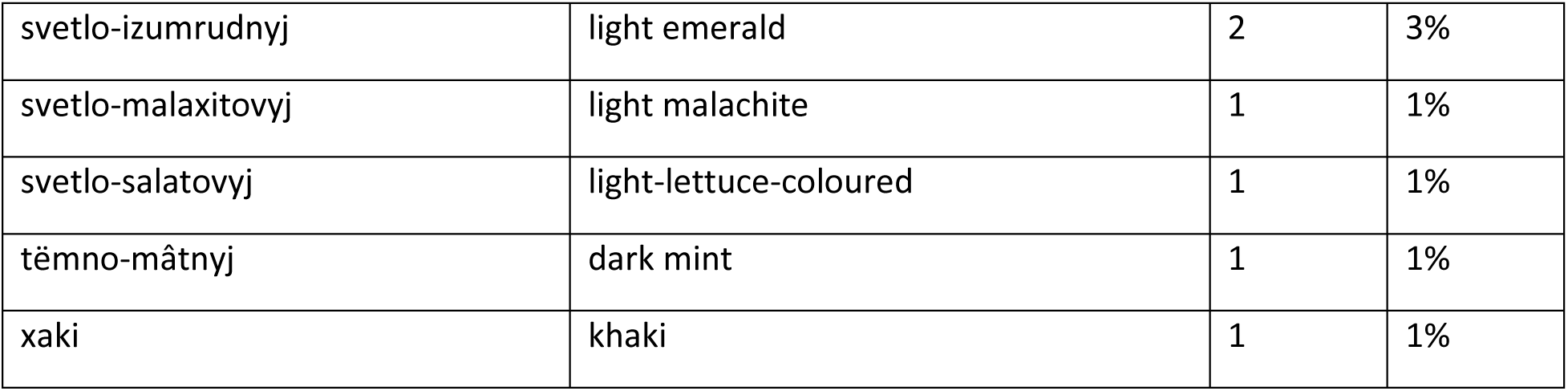
Full list (N=17) of Russian colour terms elicited for the light green stimulus with English glosses. Each term is provided with its absolute frequency (F) and relative frequency (%F), in descending order. Colour descriptors containing *zelënyj* ‘green’ are shaded.

| Colour name | English gloss | F | %F |
| --- | --- | --- | --- |
| svetlo-zelënyj | light green | 24 | 32% |
| zelënyj | green | 22 | 29% |
| mâtnyj | mint | 13 | 17% |
| salatovyj | lettuce-coloured | 2 | 3% |
| zelëno-birûzovyj | turquoise green | 2 | 3% |
| bledno-birûzovyj | pale turquoise | 1 | 1% |
| bledno-izumrudnyj | pale emerald | 1 | 1% |
| bledno-zelënyj | pale green | 1 | 1% |
| cvet morskoj volny | colour of sea wave | 1 | 1% |
| izumrudnyj | emerald | 1 | 1% |
| malaxitovyj | malachite | 1 | 1% |
| svetlo-birûzovyj | light turquoise | 1 | 1% |
| svetlo-izumrudnyj | light emerald | 2 | 3% |
| svetlo-malaxitovyj | light malachite | 1 | 1% |
| svetlo-salatovyj | light-lettuce-coloured | 1 | 1% |
| tëmno-mâtnyj | dark mint | 1 | 1% |
| xaki | khaki | 1 | 1% |

A similar picture emerged for dark green, which was named *tëmno-zelënyj* ‘dark green’ or *zelënyj* ‘green’ by 71% participants, while another 5% of participants used the term *zelënyj* ‘green’ as part of compounds *bolotno-zelënyj* ‘marsh green’, *izumrudno-zelënyj* ‘emerald green’, *malaxitovo-zelënyj* ‘malachite green’, *nasyŝennyj tëmno-zelënyj* ‘saturated dark green’, *zelënyj kak trava* ‘grass green’ (Table S2).

**Table S2.** Full list (N=13) of Russian colour terms elicited for the dark green stimulus with English glosses. Each term is provided with its absolute frequency (F) and relative frequency (%F), in descending order. Colour descriptors containing *zelënyj* ‘green’ are shaded.

| Colour name | English gloss | F | %F |
| --- | --- | --- | --- |
| tëmno-zelënyj | dark green | 39 | 51% |
| zelënyj | green | 15 | 20% |
| izumrudnyj | emerald | 9 | 12% |
| bolotnyj | marsh | 3 | 4% |
| tëmno-izumrudnyj | dark emerald | 2 | 3% |
| bolotno-travânoj | marsh grass green | 1 | 1% |
| bolotno-zelënyj | marsh green | 1 | 1% |
| izumrudno-zelënyj | emerald green | 1 | 1% |
| kamuflâžnyj | camouflage | 1 | 1% |
| malaxitovo-zelënyj | malachite green | 1 | 1% |
| nasyšennyj tëmno-zelënyj | saturated dark green | 1 | 1% |
| xvojnyj | conifer | 1 | 1% |
| zelënyj kak trava | grass green | 1 | 1% |

The light blue stimulus was labelled *goluboj* by 32% of observers, while other 42% used different compounds containing *goluboj*: *svetlo-goluboj* ‘light *goluboj*’ (12%), *nebesno-goluboj* ‘sky *goluboj*’ (12%), *pastel’no-goluboj* ‘pastel *goluboj’* (7%), *sero-goluboj* ‘grey-*goluboj’* (3%), *blednyj nebesno-goluboj,* ‘pale sky *goluboj’*, *bleklo-goluboj* ‘faded *goluboj’*, *cvet golubyx džins* ‘colour of *goluboj* jeans’, *golubo-lilovyj* ‘*goluboj*-mauve’, *golubo-seryj* ‘*goluboj*-grey’, *golubovato-sirenevyj* ‘*goluboj*-ish-lilac’, *sirenevo-goluboj* ‘lilac-*goluboj’*, *tëmno-goluboj* ‘dark *goluboj’* (1% each) (Table S3).

**Table S3.** Full list (N=21) of Russian colour terms elicited for the light blue stimulus with English glosses. Each term is provided with its absolute frequency (F) and relative frequency (%F), in descending order. In compound and modified terms containing *goluboj* and *sinij*, the original Russian form is retained in the gloss to indicate their denotative distinction. Colour descriptors containing *goluboj* are shaded.

| Colour name | English gloss | F | %F |
| --- | --- | --- | --- |
| goluboj | *goluboj* | 24 | 32% |
| svetlo-goluboj | light *goluboj* | 9 | 12% |
| nebesno-goluboj | sky *goluboj* | 9 | 12% |
| nebesnyj | sky-coloured | 8 | 11% |
| pastel'no-goluboj | pastel *goluboj* | 5 | 7% |
| svetlo-sinij | light *sinij* | 3 | 4% |
| sero-goluboj | grey-*goluboj* | 2 | 3% |
| lazurnyj | azure | 2 | 3% |
| vasil'kovyj | cornflower blue | 2 | 3% |
| blednyj nebesno-goluboj | pale sky *goluboj* | 1 | 1% |
| bleklo-goluboj | faded *goluboj* | 1 | 1% |
| cvet golubyx džins | colour of *goluboj* jeans | 1 | 1% |
| golubo-lilovyj | *goluboj*-mauve | 1 | 1% |
| golubo-seryj | *goluboj*-grey | 1 | 1% |
| golubovato-sirenevyj | *goluboj*-ish-lilac | 1 | 1% |
| lavandovyj | lavender | 1 | 1% |
| ledânoj | icy | 1 | 1% |
| nalët so slivy | plum bloom | 1 | 1% |
| sirenevo-goluboj | *lilac-goluboj* | 1 | 1% |
| svetlyj džinsovyj | light denim | 1 | 1% |
| tëmno-goluboj | dark *goluboj* | 1 | 1% |

Dark blue stimulus was named as *sinij* or *tëmno-sinij* ‘dark *sinij’* by 81% participants, with other 12% opting for *sinij* as a part of compounds: *bledno-sinij* ‘pale *sinij’*, *kosmičeskij sinij* ‘space *sinij’* (3% each), *korolevskij sinij* ‘royal *sinij’*, *priglušennyj tëmno-sinij* ‘muted dark *sinij’*, *prusskij sinij* ‘Prussian *sinij’*, *pyl’no-sinij* ‘dusty *sinij’*, *sero-sinij* ‘grey-*sinij’*, *sine-seryj* ‘*sinij*-grey’ (1% each) (Table S4).

**Table S4.** Full list (N=14) of Russian colour terms (N=14) elicited for the dark blue stimulus with English glosses. Each term is provided with its absolute frequency (F) and relative frequency (%F), in descending order. In compound and modified terms containing *sinij*, the original Russian form is retained in the gloss to indicate its denotative distinction from *goluboj*. Colour descriptors containing *sinij* are shaded.

| Colour name | English gloss | F | %F |
| --- | --- | --- | --- |
| tëmno-sinij | dark *sinij* | 33 | 43% |
| sinij | *sinij* | 29 | 38% |
| bledno-sinij | pale *sinij* | 2 | 3% |
| kosmičeskij sinij | space *sinij* | 2 | 3% |
| černičnyj | blueberry coloured | 1 | 1% |
| kobal'tovyj | cobalt | 1 | 1% |
| korolevskij sinij | royal *sinij* | 1 | 1% |
| nočnoe nebo | night sky | 1 | 1% |
| priglušennyj tëmno-sinij | muted dark *sinij* | 1 | 1% |
| prusskij sinij | Prussian *sinij* | 1 | 1% |
| pyl'no-sinij | dusty *sinij* | 1 | 1% |
| sero-sinij | grey-*sinij* | 1 | 1% |
| sine-seryj | *sinij*-grey | 1 | 1% |
| tëmno-fioletovyj | dark purple | 1 | 1% |

### Supplementary Material 3 – Differences between ‘warm’ and ‘cool’ colours, which differ in basic colour terms across light and dark examples (Experiment 2)

#### Best fitting linear mixed effect model

Fixed effects of deviancy (standard – reference level, or deviant), colour (cool – reference level, or warm) and lightness (dark – reference level, or light) and random by-participant intercepts only.

Model: amplitude ∼ deviancy*lightness*colour + 1|participant

| Predictors | Amplitude (μV) |  |  |
| --- | --- | --- | --- |
|  | Estimates | 95% CI | P |
| (Intercept) | 1.56 | [0.83,2.28] | <0.001 |
| Deviancy | -0.20 | [(-0.38),(-0.02)] | 0.029 |
| Colour | -0.04 | [(-0.21),0.14] | 0.689 |
| Lightness | -0.50 | [(-0.68),(-0.32)] | <0.001 |
| Deviancy × Colour | 0.11 | [(-0.24),0.47] | 0.525 |
| Deviancy × Lightness | -0.24 | [(-0.59),0.12] | 0.193 |
| Colour × Lightness | 0.87 | [0.51, 1.22] | <0.001 |
| Deviancy × Colour × Lightness | 1.56 | [0.85,2.27] | <0.001 |
| Random Effects |  |  |  |
| $\sigma^2$ | 1.14 | | |
| $\tau_{00}$ participant | 4.86 | | |
| ICC | 0.81 |  |  |
| $N_{\text{participant}}$ | 36 | | |
| Observations | 562 |  |  |
| Marginal $R^2$ / Conditional $R^2$ | 0.026 / 0.814 | | |

Prior to statistical analyses, we rejected any datapoints that were outliers: this resulted in rejection of 14 datapoints (as can be seen from Observations row in the table above).

Assumption checks were performed using the DHARMa R package and these confirmed that the data were appropriate for the fitted model. We also verified that the predictors were not correlated by fitting an additive model and evaluating it with the *check_collinearity* function from the performance R package.

The best fitting model is identical to the full model we initially fitted, as there was a significant 3-way interaction between colour, lightness and deviancy (F(1,518.58)=18.506, p<.001). Figure S3 depicts this interaction, with Bayesian stats pictorially coded by the dot size (smallest dot – evidence of absence, BF<0.33; medium dot –moderate evidence for a significant difference (3 < BF < 10); large dot – strong evidence of a deflection in the difference wave, BF > 10):

**Figure S3.**
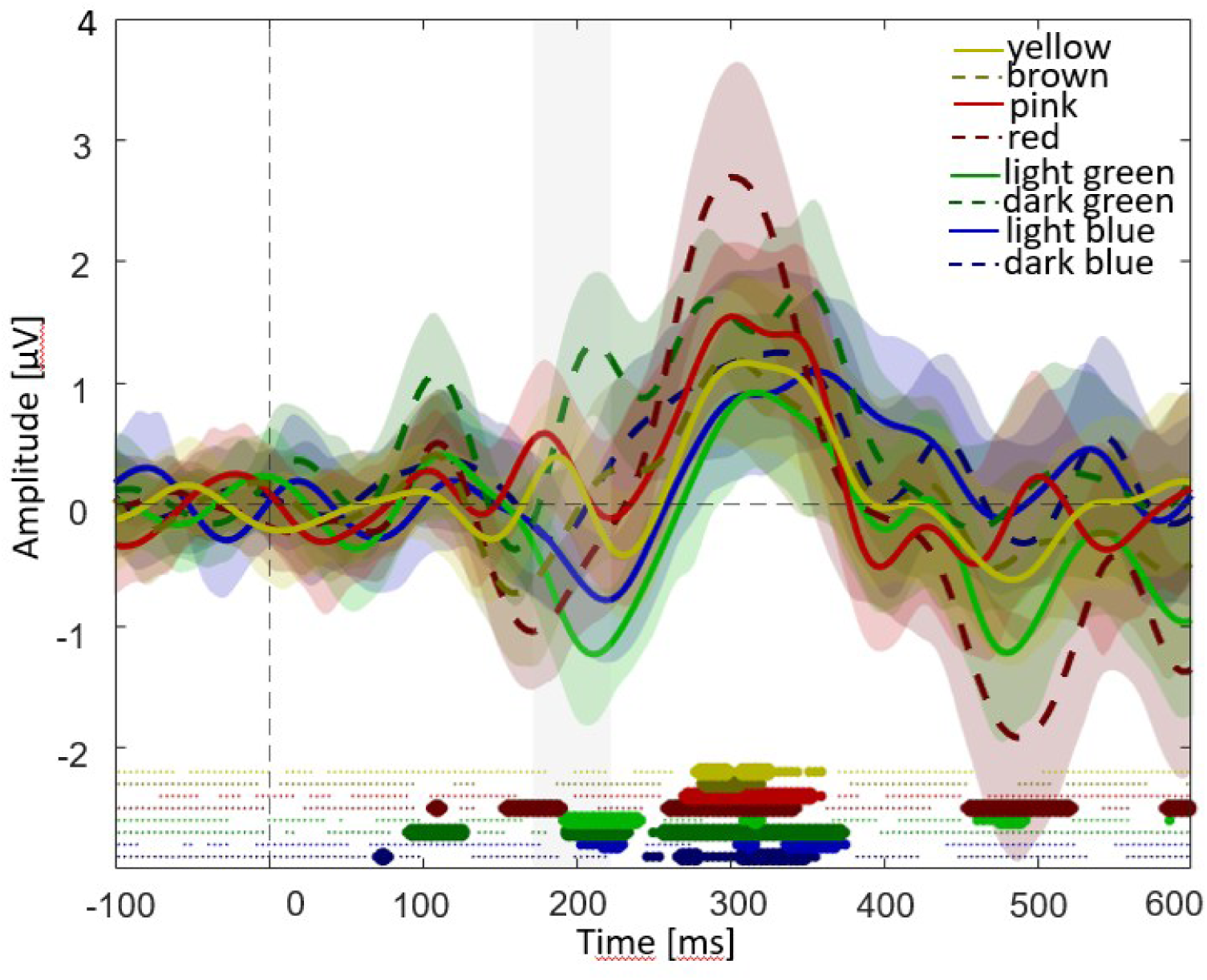
Difference waves elicited by dark and light versions of each colour, with the vMMN window indicated by a shaded grey area around 200 ms. Colour-shaded areas around each wave indicate the 95% between-subject CIs. As in the plots in the main manuscript, Bayes factor values are indicated above the x axis.

The plot reveals that the three-way interaction between colour, lightness and deviancy is likely to be driven by larger negative deflections for lighter and cooler colours. The difference wave for red also appears to be significantly more negative, but in a slightly earlier time-window, while the brown, yellow and pink difference waves did not exhibit an early negativity.

Finally, we also analysed if the difference waves depended on the order of the standard in the stimulus train – as the non-target deviant always followed a stream of 3, 4 or 5 standards. This is depicted in Figure S4 below.

**Figure S4.**
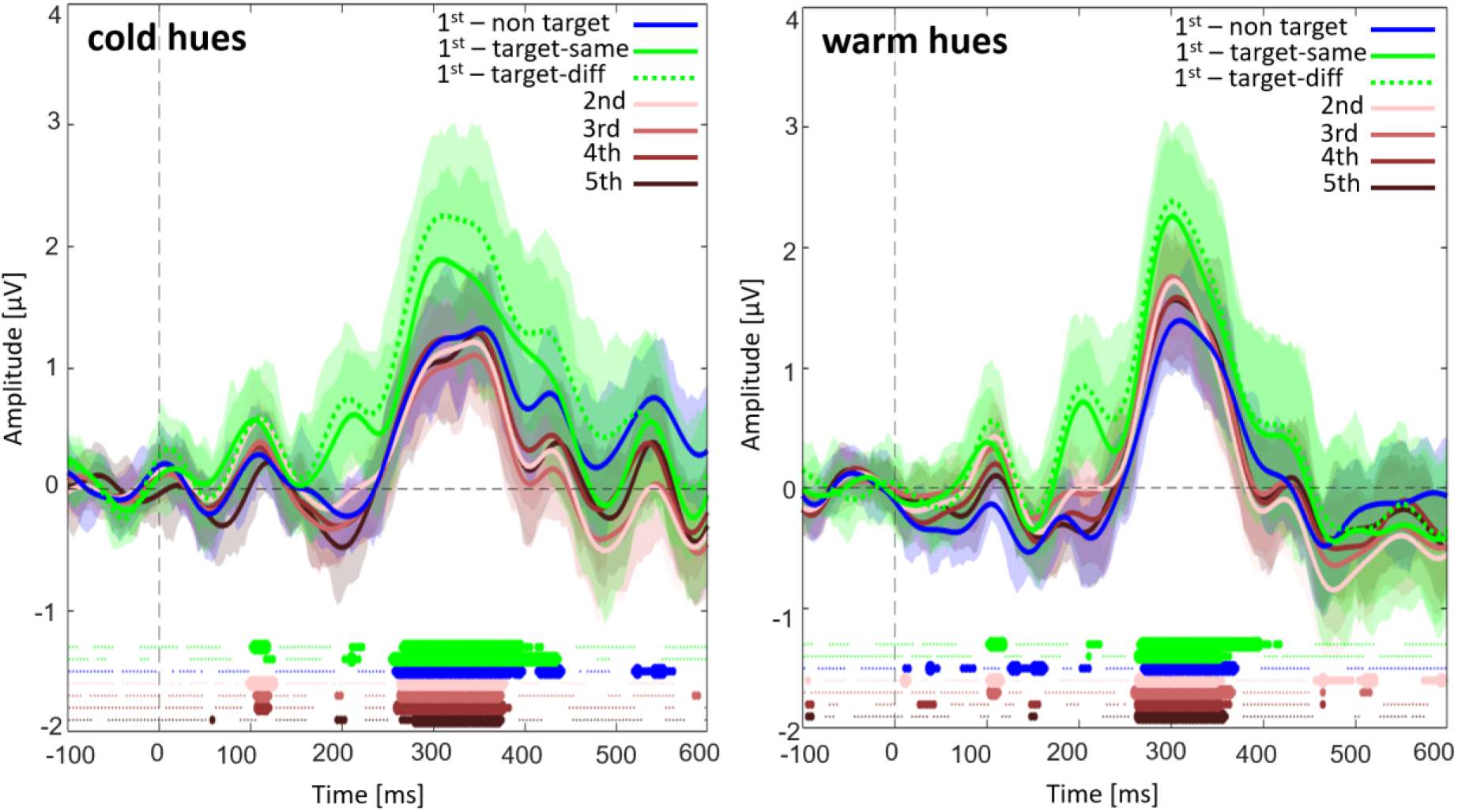
Difference waves for cool (left) and warm (right) hues calculated separately for each standard (1^st^-5^th^, with initial standards also separated depending on the preceding deviant, so that 1^st^ -non-target is the 1^st^ standard preceded by a non-target deviant, 1^st^ target-same is the first standard preceded by a same colour target, and 1^st^ target-different is the 1^st^ standard preceded by a different colour target). For both types of hues, responses significantly different to baseline in the P1 window are elicited by 1^st^ standards that follow target stimuli (i.e., squares) and 2^nd^ standards – in parallel, strongest negative responses in the N1 window are observed for the 5^th^ standards. Furthermore, 1^st^ standards that follow a target elicit a positive peak in the difference wave in the same time-window as the vMMN. This is likely the reason why trials following a target are often excluded from vMMN analysis (e.g., Male et al., 2020).

#### Power analysis

Sensitivity analyses on the best fitting model were performed by resampling the number of participants (1000 simulations) and evaluating the observed power for obtaining the statistically significant three-way interaction, using the R package *simr*. We evaluated the power to detect the observed 3-way interaction if effect sizes were reduced by 15%, as advised by Kumle and colleagues (2021), and found that this was 68.50% (95% CI [65.52%, 71.37%]); for effect sizes reduced by 10%, we had 76.10% power (95% CI [73.33, 78.71%]) and for effect size reduced by 5% we had 79.20% power (95% CI [76.55, 81.68%]).

Power simulations therefore show that we have just about enough power to observe a three-way within-subject interaction conceptually similar to the interaction observed in Thierry et al. (2009). We had 36 participants, while their sample size was 20. The reported partial eta squared (μ_p_^2^ = 0.112) for the mixed-effect (i.e., between-/within-participants) three-way interaction in Thierry et al. (2009) was associated with a 95% confidence interval whose lower bound was close to zero (0.004-0.273), which indicates imprecision and suggests that the effect they observed may range from very small to moderately large.

#### Free colour naming of colour circles by English speaking participants

1) **light blue:** blue n=21, light blue n=8, sky blue n=3, baby blue n=2, pale blue n=1, periwinkle n=1;
2) **dark blue:** blue n=29, dark blue n=4, deep blue n=1, royal blue n=1, turquoise n=1;
3) **light green:** green n=16, light green n=7, mint n=5, turquoise n=3, aquamarine n=1, lime n=1, matcha green n=1, sage green n=1, sage n=1;
4) **dark green:** green n=28, dark green n=5, forest green n=1, olive green n=1, olive n=1;
5) **pink:** pink n=27, light pink n=5, red n=2, rose n=1, salmon pink n=1;
6) **red:** pink n=10, red n=8, dark pink n=4, maroon n=4, magenta n=2, rose n=1, raspberry n=1, burgundy n=1, mauve n=1, purple n=1, purpley pink n=1, brown-red n=1, pinky maroon n=1;
7) **yellow:** yellow n=30, light yellow n=3, pale yellow n=2, green n=1;
8) **brown:** green n=11, brown n=11, dark green n=4, olive green n=2, olive n=1, hazel n=1, swamp n=1, taupe n=1, yellow n=1, khaki n=1.

### Supplementary Material 4 – Early negativities driven by differences in hue, saturation and luminance contrast (Experiment 3)

Initially, we fitted a model with the full set of fixed effects (deviancy*colour*lightness*saturation), but the step function from lmerTest R package indicated that hue (with two levels: red and green) contributed significantly to the model only on its own, rather than in interaction with the other three factors, which also failed to interact in a 3-way fashion with each other:

Deviancy:Hue:Saturation:Lightness, F(1, 654.99) = 0.0174, p = 0.895
Deviancy:Hue:Saturation, F(1, 655.99) = 0.0198, p = 0.888
Hue:Saturation:Lightness, F(1, 656.99) = 0.2164, p = 0.642
Hue:Saturation, F(1, 657.99) = 0.8408, p = 0.359
Deviancy:Hue:Lightness, F(1, 658.99) = 1.1350, p = 0.287
Deviancy:Hue, F(1, 659.99) = 0.1456, p = 0.703
Hue:Lightness, F(1, 660.99) = 0.6421, p = 0.423
Deviancy:Saturation:Lightness, F(1, 662.00) = 3.091, p = 0.079

A series of two-way interactions remained in the model, together with the fixed effect of hue:

Hue, F(1, 663.00) = 9.706, p = 0.002**
Deviancy:Saturation, F(1, 663.00) = 7.301, p = 0.007**
Deviancy:Lightness, F(1, 659.99) = 4.994, p = 0.026*
Saturation:Lightness, F(1, 660.99) = 6.427, p = 0.011*

The best-fitting model properties are listed below:

amplitude ∼ deviancy + hue + saturation + lightness + (1 | participant) + deviancy:saturation + deviancy:lightness + saturation:lightness

| Predictors | Amplitude ($\mu V$) | | |
| --- | --- | --- | --- |
|  | Estimates | 95% CI | P |
| (Intercept) | 1.94 | [1.17, 2.71] | **<0.001** |
| Deviancy | -0.35 | [(-0.56), (-0.14)] | **<0.001** |
| Hue | 0.28 | [0.10, 0.46] | **0.002** |
| Saturation | -0.17 | [(-0.39), 0.04] | 0.109 |
| Lightness | -0.23 | [(-0.45), (-0.02)] | **0.031** |
| Deviancy × Saturation | 0.57 | [0.16, 0.98] | **0.007** |
| Deviancy × Lightness | -0.47 | [(-0.88), (-0.06)] | **0.026** |
| Saturation ×<br>Lightness | -0.55 | [(-0.97), (-0.12)] | **0.011** |
| Random Effects |  |  |  |
| $\sigma^2$ | 1.43 | | |
| $\tau_{00}$ participant | 3.75 | | |
| ICC | 0.72 |  |  |
| $N_{\text{participant}}$ | 25 | | |
| Observations | 695 |  |  |
| Marginal $R^2$ / Conditional $R^2$ | 0.019 / 0.729 | | |

Post-hoc tests via emmeans package (using multivariate t distribution method for correction for multiple comparisons) revealed the following drivers for the observed interactions and the main effect of hue:

#### For saturation x lightness

Lower lightness was associated with lower amplitudes only when the stimulus was also desaturated (all other ps > 0.872):

higher saturation and lightness vs. desaturated lower lightness: 0.407, 95%CI [0.143, 0.672], t(663)= 3.015, p = 0.0139

desaturated of higher lightness vs. desaturated lower lightness: 0.508, 95%CI [0.175, 0.841], t(663)= 2.985, p = 0.0151

higher saturation and lower lightness - desaturated lower lightness: 0.4482, 95%CI [0.115, 0.7814], t(663) = 2.634, p = 0.0413.

#### For saturation x deviancy

Higher saturation standards elicited higher amplitudes (all other ps > 0.635):

Higher saturation standard vs. higher saturation deviant: 0.6363, 95%CI [0.393, 0.879], t(663) = 5.116, p <.0001

Higher saturation standard vs. desaturated standard: 0.4580, 95%CI [0.162, 0.754], t(663) = 3.043, p = 0.0128

Higher saturation standard vs. desaturated deviant: 0.5258, 95%CI [0.228, -0.824], t(663) = 3.464, p = 0.0030

#### For lightness x deviancy

Lower lightness deviants elicited lower amplitudes (all other ps > 0.781):

standard higher lightness - deviant lower lightness: 0.58541, 95%CI [0.287, 0.883], t(663) = 3.856, p< 0.001

deviant higher lightness - deviant lower lightness: 0.46846, 95% CI [0.173, 0.764], t(663) = 3.095, p = 0.0107

standard lower lightness - deviant lower lightness: 0.58713, 95% CI [0.254, 0.920], t(663) = 3.451, p = 0.0034

#### For hue

Green elicits lower amplitude than red:

green vs. red: -0.283, 95%CI [(−0.105), (−0.461)], t(663) = -3.115, p = 0.0019

Finally, we tested how stimuli with lower lightness or saturation sum up to evaluate the degree to which lightness and saturation are processed by separable neural resources. If this were the case, the predicted waveform for feature conjunctions (higher lightness and saturation or lower lightness and desaturated) should correspond to predictions made by averaging waveforms for single feature changes. Figure S5 below depicts predicted (full line) and observed (dotted line with grey 95% CIs) waveforms.

**Figure S5.**
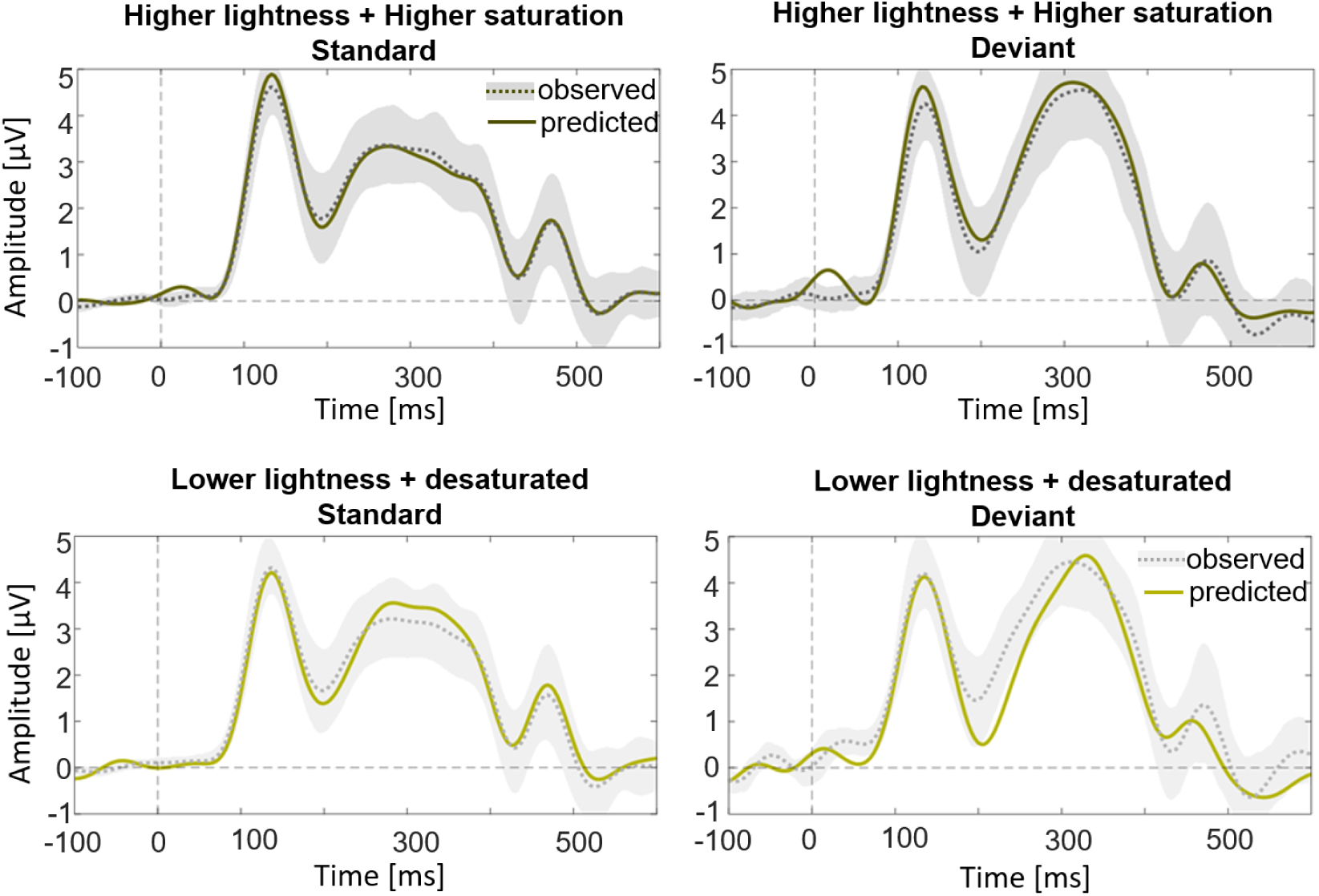
Visual evoked potentials (VEPs) observed in conditions that combined lightness and saturation changes, as well as waveforms predicted by averaging responses from VEPs elicited by changes in only one of the two features (lightness or saturation). Shaded grey area corresponds to a 95% between-subject CI for observed data.

As can be seen, the departure from the predicted waveform can be seen during the N1—P300 transition in the desaturated, lower lightness deviants. These stimuli were recorded in blocks with higher and more saturated standards, while single features, from which the predictions are derived, come from blocks in which the standards the participants were adapted to possessed only one of these two features. Observed responses to these stimuli in the N1—P300 window show amplitude that is reduced relative to what we predicted from single feature blocks. This is in line with shared neural resources between colour and luminance processing, which therefore can adapt more significantly (exhibiting repetition suppression).

